# Direct single-molecule detection and super-resolution imaging with a low-cost portable smartphone-based microscope

**DOI:** 10.1101/2024.05.08.593103

**Authors:** Morgane Loretan, Mariano Barella, Nathan Fuchs, Samet Kocabey, Karol Kołątaj, Fernando D. Stefani, Guillermo P. Acuna

## Abstract

We present a novel, low-cost, portable smartphone-based fluorescence microscope capable of directly detecting single molecules without signal amplification. The setup leverages the image sensors and data handling capacity of mass-produced smartphones, making it adaptable to any smartphone and capable of detecting single molecules across the visible spectral range. We showcase this capability through single-molecule measurements on DNA origami models and super-resolution microscopy of biological cells by single-molecule localization microscopy. This development paves the way for biotechnology innovations making use of massively distributed or personalized assays with single-molecule sensitivity with the potential to revolutionize digital bioassays, point-of-care diagnostics, field expeditions, STEM outreach, and life science education.

---

Presently, the global proliferation of smartphones is estimated to be around 7 billion units^1^. This vast scale of production continually propels the integration of technological advancements at a cost-efficiency unparalleled by other sectors. The unrivaled portability, compactness, and worldwide accessibility, coupled with high-performance image sensors, robust computing power, and connectivity, have catalyzed the development of specialized smartphone-based setups for Point-Of-Care (POC) and personal biomedical applications^2^. These setups leverage the advanced camera technology of modern smartphones^3^ for fluorescence-based clinical diagnostics^4^, quantification of immunoassays^5,6^, detection of bacteria^7^, cancer cytology^8^, fresh tissue imaging^9^, and environmentally significant measurements such as lead and microplastics quantification^10,11^. Naturally, there has been a quest to take these assays to the single-molecule level. Smartphone-based portable setups with single-molecule sensitivity would unlock all the advantages of sub-ensemble determinations for personal or POC devices. For instance, it would be possible to quantify target molecules under ultra-low analyte concentrations by implementing digital bioassays^12,13^ or to conduct super-resolution fluorescence imaging^14^.

Since the pioneering work by the Ozcan group a decade ago^15^, which detected fluorescent beads, the sensitivity of smartphone-based fluorescence microscopes has been rigorously benchmarked and continually improved^16^. However, the ultimate goal of detecting single fluorescent molecules has remained a challenge. Until now, detecting single or a few fluorescent molecules in smartphone-based setups was only possible using physical or chemical amplification mechanisms, which are hardly compatible with large-scale point-of-care or personal applications. For instance, single-molecule fluorescence was detected using a smartphone microscope by strongly enhancing the fluorescence signal with DNA origami-based optical antennas^17^. Alternatively, chemical amplification approaches have been implemented to increase the number of copies of the analyte. Via this approach, single gene expressions were detected in a low number of HepG2 cells using a smartphone-based setup with an integrated thermal cycler for polymerase chain reaction amplification^18^. Still, to date, the direct detection of single fluorescent molecules has remained within the realm of high-end devices.

Here, we present a portable and inexpensive smartphone-based fluorescence microscope capable of directly detecting single molecules without the need for signal amplification. To assess its performance, we employed DNA origami fluorescence standards and analyzed single-molecule intensity fluctuations using three commercially available smartphones from different manufacturers. Our findings demonstrate robust single-molecule detection with a favorable signal-to-noise ratio of 3.3. Furthermore, we successfully leveraged the microscope’s single-molecule sensitivity to achieve super-resolution microscopy. By implementing DNA PAINT Single-Molecule Localization Microscopy (SMLM) on the smartphone platform, we were able to image both DNA origami structures and microtubule networks. The acquired images exhibited a localization precision of 86 nm, translating to an 11-fold enhancement in resolution. In summary, this work represents a significant advancement towards making single-molecule fluorescence assays and methods universally accessible. This novel smartphone-based microscope has the potential to revolutionize various fields, from point-of-care quantitative sensing devices to nanoscale imaging for personalized diagnostics.

## Results

### The smartphone-based microscope

Figure 1a includes a photograph of the smartphone-based fluorescence microscope. The microscope weighs 1.2 kg, and its dimensions are 11 cm × 22 cm × 12 cm, smaller than a standard shoe box. It is a stand-alone unit that includes a laser, a power source, and the necessary optical components in a modular design conceived to optimize sensitivity, portability, affordability, and practicality (additional photographs can be found in Supplementary Information S1). It can be operated on any conventional table or on the floor and requires minimal user training and standard sample preparation techniques. The cost of the microscope lies under 350 € (see list of components and respective price in Methods and Supplementary Information S2). Finally, and as opposed to other designs that focus on a particular model, our microscope is flexible enough to host any smartphone with a camera, although, for this work, we benchmarked its performance with flagship models from the three main global smartphone producers, Apple, Samsung, and Huawei.

**Figure 1:**
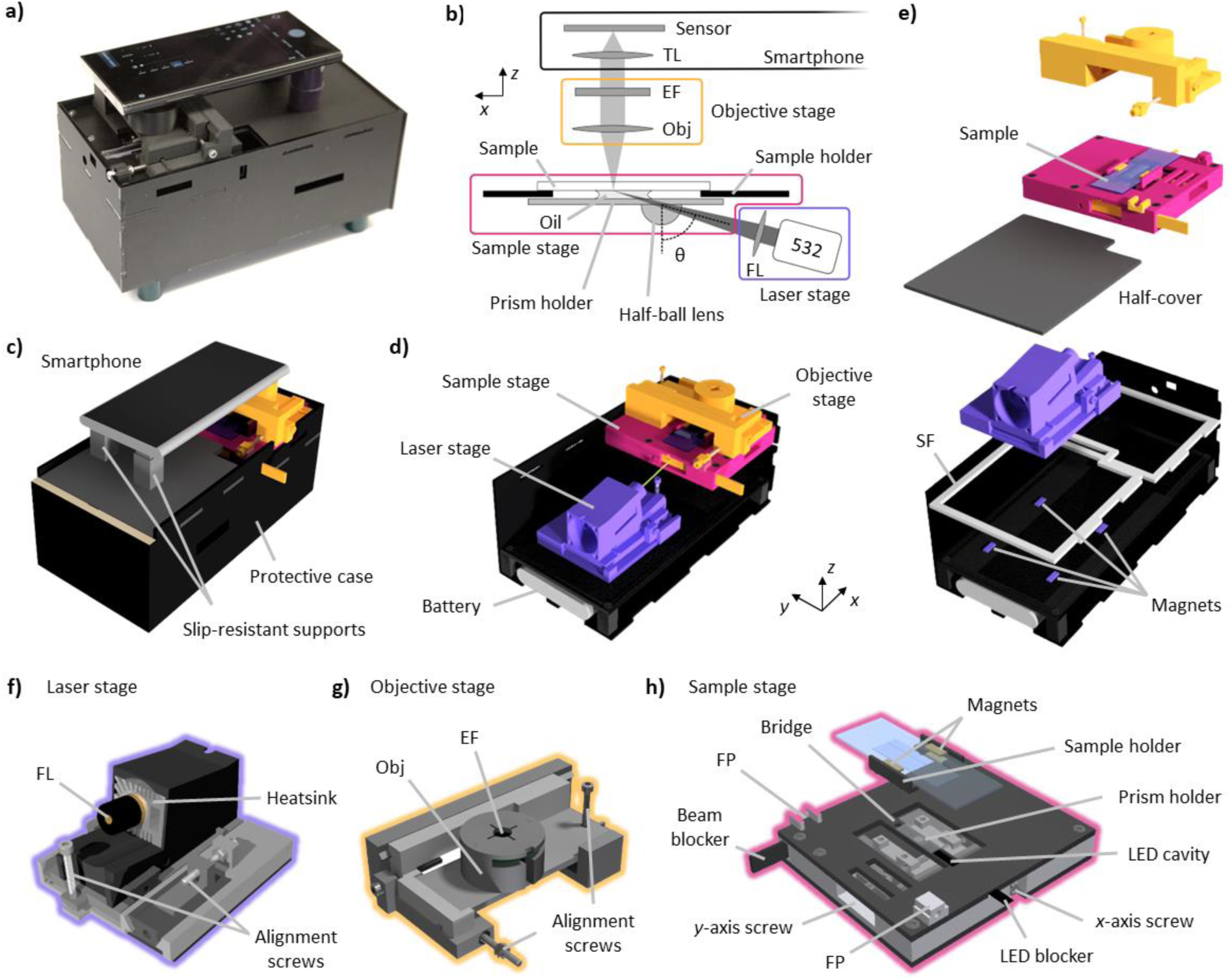
Smartphone-based fluorescence microscope. (a) Photograph of the setup with a Samsung Galaxy S22 Ultra on top. (b) Optical path sketch of the smartphone-based microscope. Each color frame encloses the optical components of the objective stage (yellow), sample stage (pink), laser stage (purple), and the smartphone camera (black). (c) Rendered model of the smartphone-based microscope ready for use. (d) Inside view with the laser, sample, and objective stages. (e) Exploded view depicting the supporting frame (SF) and the laser stage magnets. (f-h) Details of the three stages of the microscope with their main components TL: tube lens. FL: focusing lens. EF: emission filter. FP: fixation point.

Figure 1b depicts a sketch of the optical paths introduced to enhance the microscope’s sensitivity by minimizing the background signal. Fluorescence excitation is achieved with a laser beam that passes through a focusing lens (FL) and reaches a half-ball lens that acts as a prism for total internal reflection (TIR) illumination. The half-ball is glued to a glass slide (prism holder) using an optical adhesive. Immersion oil is applied between the prism holder and the sample substrate to match the refractive indexes and complete the TIR configuration. This configuration differs from previous smartphone-based microscopes that used waveguided LEDs as light sources, which, in contrast to lasers, are less radiant and spectrally broader^16,17,19–25^. The light emitted by the sample is collected by an inexpensive, low numerical aperture (NA) air objective (Obj), spectrally selected with an emission filter (EF), and focused onto the smartphone CMOS sensor by its camera lens, acting as a tube lens (TL).

Figure 1b also depicts the modular design of the smartphone-based microscope. The setup is composed of four parts: the protective black case, the laser stage (purple), the objective stage (yellow), and the sample stage (pink). The protective black case counts with a top half-cover that shields the user from laser radiation and allows the smartphone to be placed using two slip-resistant silicone supports that can be freely moved. This simple configuration makes the setup compatible with smartphones of different sizes and camera positions (Figure 1c). The protective case also hosts the laser control electronics and the battery, as shown in Figure 1d. During transportation, it is used to store all the optical components, immersion oil, and samples (see Supplementary Information S1). Inside, four magnets allow the laser stage to be easily removed and repositioned without needing beam realignment (Figure 1e). This gives the setup flexibility for selecting the excitation wavelength, as different lasers can be easily interchanged. The removable sample and objective stages fit into a supporting frame (SF, Figure 1e).

The laser stage shown in Figure 1f comprises the laser module with the focusing lens, a heatsink, and alignment screws. A cooling fan at the end of the heatsink is optionally mounted for high-power lasers. The fan is turned off or removed to reduce possible vibrations when high localization precision is desired, like during SMLM measurements. The laser stage counts with three degrees of freedom that allow translation over the microscope horizontal plane (*xy*) and fine-tuning the incidence angle (*θ*) to achieve highly inclined and laminated optical sheet (HILO) or TIR illumination.

The objective stage shown in Figure 1g has been conceived to focus on the sample plane (*z*-axis) and to align the objective with the illuminated area of the sample (*xy* plane). These degrees of freedom, controlled by alignment screws, offer flexibility when using different sample substrates without the need to realign the laser. The objective holder contains the low-cost objective and the emission filter. The emission filter is installed in a lateral slot, enabling easy exchange without touching the smartphone. The objective can also be exchanged if required.

To be able to move the sample over *x* and *y* directions, we designed a sample stage (Figure 1h) that houses two different moving elements operated by two screws. On the moving stage, the sample holder includes magnets to secure the sample with a top holder. The prism holder is glued to the stage on the bridge, right below the sample. A white light-emitting diode (LED) is incorporated in a cavity inside the bridge, next to the prism, that allows image pre-focusing, sample positioning, and bright-field imaging. An LED blocker is used to prevent the laser from exciting undesirably the LED. Similarly, a manual shutter (Beam blocker) is included to obstruct the laser beam from reaching the sample. Finally, the sample stage features two small fixation points (FP) that guide and fix the objective stage position.

### Direct single-molecule detection with the smartphone-based microscope

To test the ability of our portable microscope to detect the direct emission of a single fluorescent molecule, we used fluorescence standards based on DNA origami structures^26^. The design consists of a 60 × 52 nm^2^ 2-layer sheet origami (2LS) with an ATTO 542 dye at the center, and an ATTO 647N dye 22 nm to the side (Figure 2a). Six biotins are included at the bottom side for surface binding. Transmission Electron Microscopy (TEM) images of the 2LS are shown in Supplementary Information S3.

**Figure 2:**
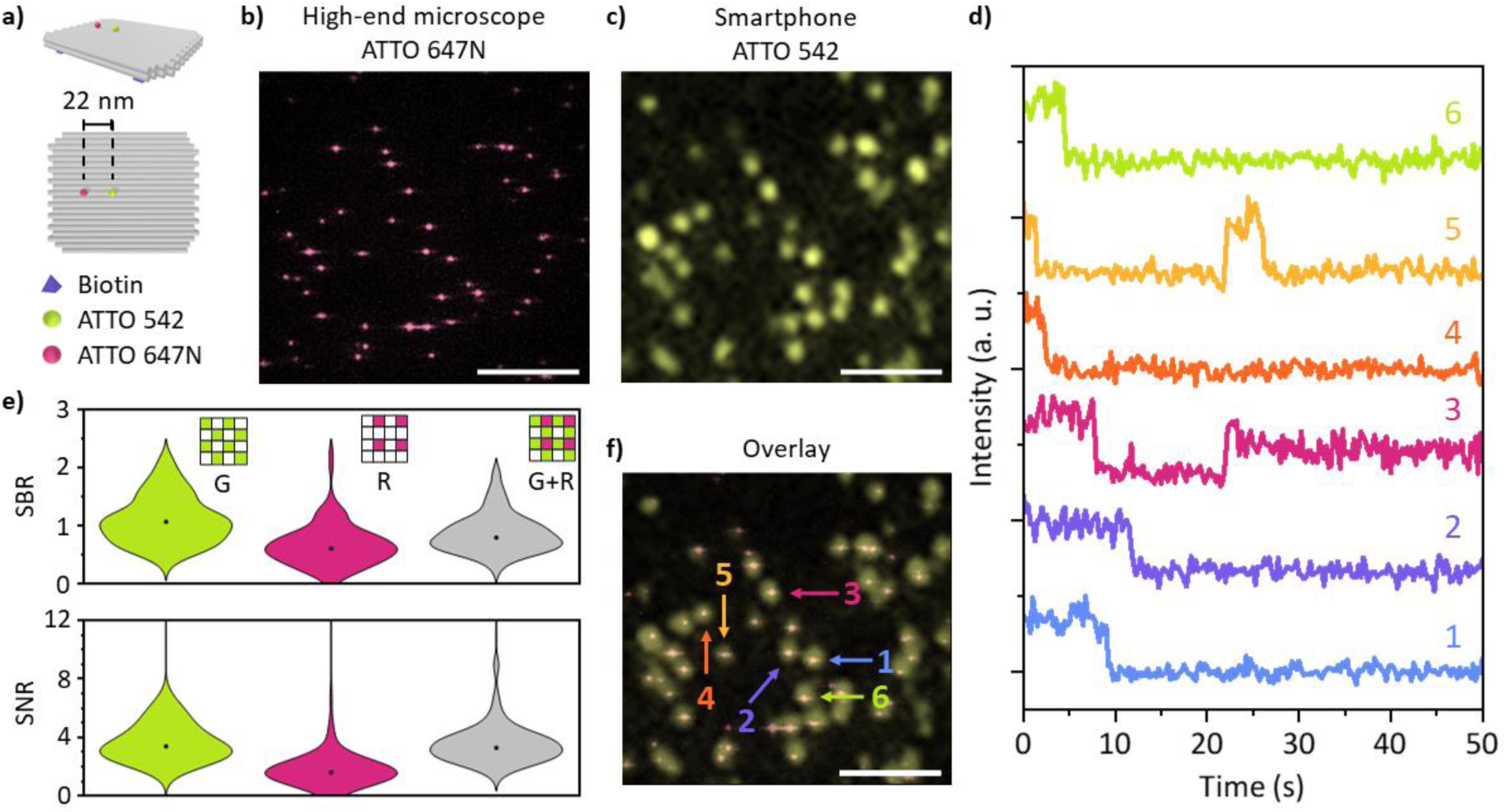
Direct single molecule detection with the smartphone-based microscope. (a) 2LS DNA origami nanostructure was used to detect single molecules with an ATTO 542 at the center and an ATTO 647N spaced 22 nm apart. Fluorescence images of the 2LS origami sample were recorded over the same area with (b) the high-end microscope detecting only the ATTO 647N and (c) the smartphone-based microscope detecting only the ATTO 542 with a Samsung Galaxy S22 Ultra. Scale bar: 10 µm. (d) Intensity traces vs. time of several ATTO 542 single molecules. They correspond to the 2LS nanostructures indicated in the overlayed image shown in (f). (e) SBR, or Weber contrast, and SNR violin plots of 104 molecules acquired with the smartphone-based microscope using the green (G) channel, the red (R) channel, and the sum of both channels, green and red (G+R). Insets: Bayer filter pixel selection. (f) Overlay of images (b) and (c). Arrows and numbers indicate intensity traces plotted in (d). Scale bar: 10 µm.

A sample with DNA origami immobilized on a quartz substrate at a density suited for single-molecule measurements was prepared. We first imaged the sample using a custom-built high-end widefield fluorescence microscope to detect the ATTO 647N dyes (see Methods). Exciting at 640 nm, we recorded the intensity over time of single ATTO 647N molecules until they bleached (see Supplementary Information S13 for experimental details). An exemplary image is included in Figure 2b where, as expected, single ATTO 647N molecules from several DNA origami 2LS can be clearly detected as a diffraction-limited spot corresponding to the point spread function (PSF) of the system. Next, the sample was moved onto the smartphone-based microscope placed over a standard desk to detect the ATTO 542 molecules excited at 532 nm. The sample position was adjusted to image the region observed previously with the high-end microscope. A Samsung Galaxy S22 Ultra smartphone was used to record the intensity over time of the ATTO 542 dyes with the MotionCam Pro app without compression (raw data mode).

Figure 2c shows the corresponding fluorescence image acquired with the smartphone-based setup in the green spectral range (see Supplementary Information S13 for experimental parameters). A first visual inspection already indicates that single ATTO 542 molecules can be detected with the smartphone-based microscope, albeit with a bigger PSF due to the lower NA of the employed objective. An overlay of the images allowed us to correlate the presence of single ATTO 542 and ATTO 647N molecules incorporated in the same 2LS DNA origami (Figure 2f) within the same diffraction-limited spot. We found that 89% of the fluorescent spots observed on the smartphone-based microscope correspond to origami nanostructures with a single ATTO 542 and a single ATTO 647N. The remaining spots belong to either origami nanostructures labeled with only a single ATTO 542 or unidentified fluorescence sources. These results are consistent with typical values for the yield of single-stranded DNA labeling with a single dye (see further details in Supplementary Information S4). This already provides solid evidence that our microscope can detect single ATTO 542 molecules. Further confirmation was obtained from fluorescence transients taken with the smartphone. Six exemplary intensity transient traces of a single ATTO 542 can be found in Figure 2d, with their corresponding positions highlighted in Figure 2f (numbered arrows). Binary blinking and single-step photobleaching events are clearly visible, which constitute unambiguous signatures of the photophysics of single fluorophores.

Finally, in order to quantify the microscopés sensitivity for single molecule detection, we estimated both the signal-to-background ratio (SBR), sometimes also referred to as Weber contrast, and the signal-to-noise ratio (SNR). Since data was acquired in raw mode, we have access to the three Bayer filter channels of the camera sensor (red, green, and blue). This enables choosing the most convenient combination of channels to maximize the SBR and the SNR depending on the target fluorophore. Figure 2e shows the SBR and the SNR distributions for 104 ATTO 542 molecules detected on the green, red, and green + red channels. The blue channel was not considered, as the long-pass emission filter used had a cut-off wavelength of 550 nm. It is worth mentioning that the SBR was estimated from one frame and, therefore, quantifies the extent to which a single molecule can be detected from an image, whereas the SNR was extracted from the fluorescence transients (see further details in Methods). Median values of SBR and SNR were higher for the green channel alone (SBR_G_ = 1.1, SNR_G_ = 3.3) than for the red channel alone (SBR_R_ = 0.60, SNR_R_ = 1.6) or the sum of green and red channels (SBR_G+R_ = 0.78, SNR_G+R_ = 3.2). Thus, all single-molecule data shown for ATTO 542 was analyzed from the green channel of the smartphone sensors.

Analogous measurements were performed with an iPhone 14 Pro and a Huawei P20 Pro (see Supplementary Information S5). The smartphone-based setup can detect single-molecule fluorescence with any of those smartphones, which indicates that this type of measurement can be performed with any state-of-the-art smartphone.

### Super-resolution benchmark with DNA origami models

Direct single-molecule fluorescence detection is a prerequisite for super-resolution imaging. Thus, among other exciting possibilities, our microscope has the potential to perform SMLM with nanometer resolution but using a mass-consumption smartphone. To demonstrate this, we designed a DNA origami nanostructure based on an 8-helix bundle (8HB) with two docking sites for DNA-PAINT^27^ separated by 256 nm, each composed of 11 or 12 docking strands, and biotins for binding to the surface, as depicted in Figure 3a. TEM images of several 8HBs are shown in Supplementary Information S3.

**Figure 3:**
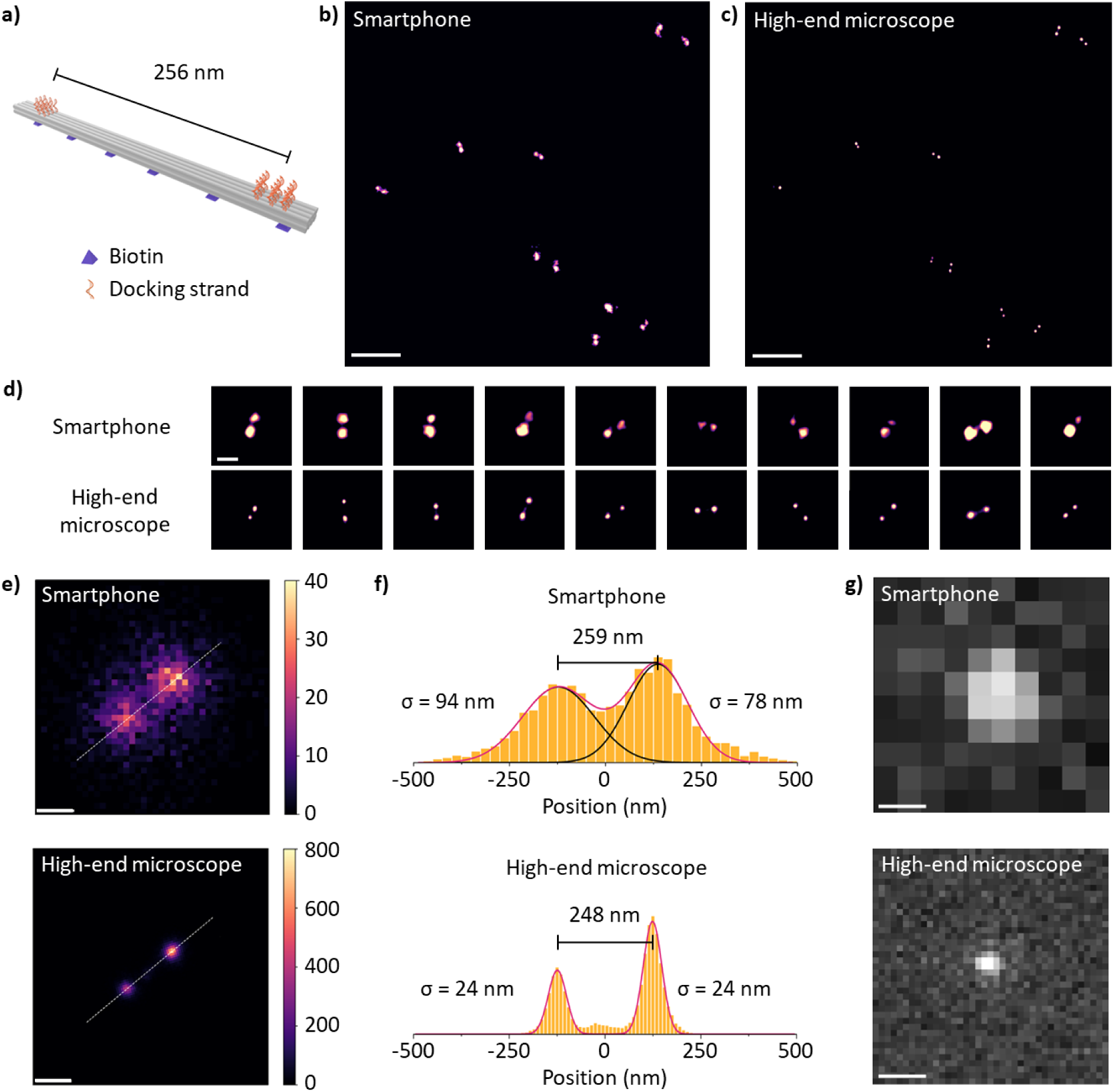
DNA-PAINT with the smartphone-based microscope. (a) 8HB DNA origami nanostructure with two docking sites and biotin surface binding functions. Super-resolved images of several nanostructures in an aqueous solution acquired with (b) the smartphone-based microscope and (c) a high-end microscope. Images are equally scaled. Scale bar: 2 µm. (d) One-to-one comparison of 10 nanostructures. Angle correspondence is observed. Scale bar: 300 nm. (e) Average 2D histograms of localizations of the same set of 10 nanostructures. Bin sizes are 25 nm for the smartphone-based setup and 10 nm for the high-end microscope. Scale bar: 150 nm. (f) Localization histograms along the central axis (dashed white line) of the nanostructure in (e). Bin size: same as in (e). (g) Typical diffraction-limited fluorescence spots, i.e., the Point Spread Function of each system, equally scaled. Scale bar: 1 µm.

Samples were prepared with 8HB DNA-origami dispersed on a quartz substrate at densities suited for their individual observation through DNA-PAINT. We used a fluorogenic imager strand with a Cy3b fluorophore on one of its ends and a BHQ-2 fluorescence quencher on the other to reduce the background level^28^. The imager had 12 complementary bases with the docking strands, delivering an average binding time of about 1 s. Gold NPs were used as fiducial markers for drift correction (further details in Methods and Supplementary Information S6).

DNA-PAINT measurements on these samples were performed both on the high-end single-molecule microscope and the smartphone-based setup. Single-molecule blinking videos were acquired and subsequently analyzed with Picasso software^29^ to reconstruct super-resolution images from single-molecule localizations. An essential step in the analysis was a 3-frame averaging of the video before analysis with Picasso. While this increased the overall SNR slightly, it improved the symmetry of the detected single-molecule signals (see Supplementary Information S6). Figures 3b and 3c show super-resolved images of DNA origami on the same region of a sample obtained from measurements on the smartphone-based setup and the high-end microscope, respectively. Remarkably, the smartphone-based setup delivers super-resolution images in excellent agreement with the high-end microscope measurements. A one-to-one comparison of individual images exhibits docking sites for DNA-PAINT with matching orientations and separation distances, as seen in the ten examples of Figure 3d. A quantitative comparison of the separation between docking sites was made by computing the average 2D histograms of localizations of those ten nanostructures (Figure 3e). Figure 3f displays the localization histogram along the long axis of the nanostructure fitted with two Gaussian functions, one for each docking site. The determined average distances between docking sites were 259 nm and 248 nm, for the measurements with the smartphone-based microscope and the high-end microscope, respectively. Both are in good agreement with the DNA origami design.

From these measurements, an estimate of the gain in resolution can be obtained from the ratio of the PSF size and the localization precision σ. Figure 3g shows exemplary PSFs for both setups. The smartphone-based microscope presents a PSF waist of (1.0 ± 0.1) µm, whereas the high-end microscope one of (0.22 ± 0.03) µm. The average localization precision with the smartphone was *σ* = 86 nm, corresponding to an increase in resolution of 11.6 times. In the case of the high-end microscope, the localization precision was *σ* = 24 nm leading to an increase in resolution of 9.2. The better value of this metric for the smartphone-based setup is most likely due to the influence of the size of the DNA-PAINT docking sites, which is in the order of 28 nm and, therefore, more comparable to the final resolution of the high-end microscope measurements. The performance of the smartphone-based setup is remarkable, especially considering that the irradiance on the sample was lower than in the high-end microscope (see Supplementary Information S13).

### Super-resolution cell imaging with the smartphone-based microscope

To definitively showcase the versatility of the smartphone-based single-molecule detection platform, we undertook super-resolution imaging of the microtubule network of U2OS cells on standard glass coverslips and using the smartphone-based setup on a common desk table. α-tubulin was immunolabeled with primary-secondary antibodies for DNA-PAINT as sketched in Figure 4a (see Methods for further details). We used Cy3b-labeled imager strands at a concentration of 500 pM, photostabilized in 1.5×PPC. Figure 4b shows a composite image displaying a super-resolved image of the microtubule network obtained with the smartphone setup and the diffraction-limited fluorescence image. These results provide compelling evidence that the smartphone-based setup can achieve super-resolution imaging using SMLM. Further, we again benchmarked its performance to the high-end single-molecule microscope. Figures 4c and 4d present super-resolved images of the same region of the microtubule network, respectively obtained from videos acquired with smartphone-based and high-end microscopes. The achieved resolution was estimated via decorrelation analysis^30–32^ yielding values of 210 nm and 66 nm, for the smartphone-based and the high-end microscopes, respectively (see further details in Supplementary Information S7). These values are in line with visual inspection of individual microtubules (Figure 4e) and the measurements on DNA origami (Figure 3).

**Figure 4:**
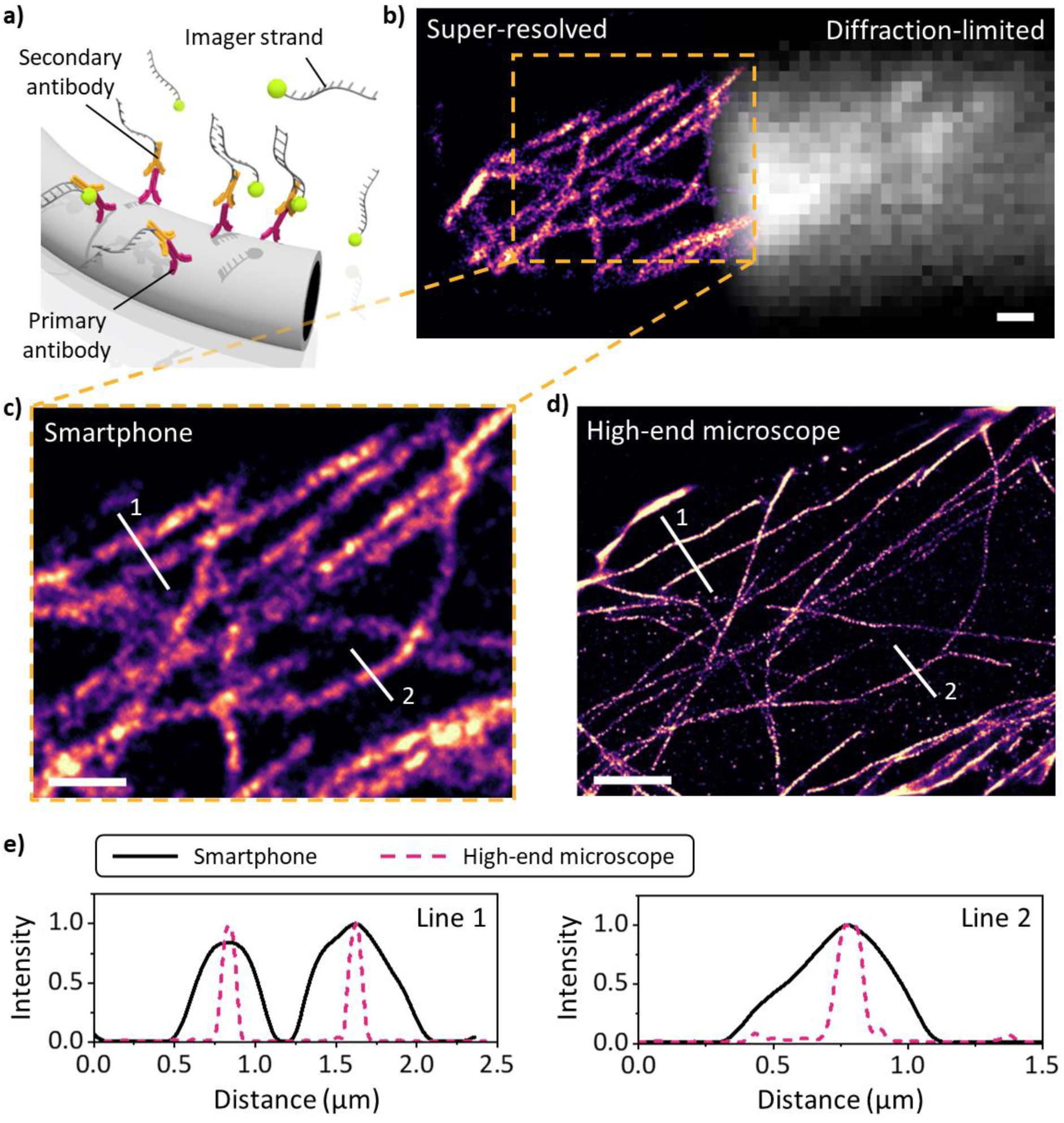
Super-resolution DNA-PAINT imaging on biological cells. (a) Sketch of an immunolabeled microtubule with primary and secondary antibodies. The secondary antibody is conjugated with a docking DNA strand complementary to the Cy3b-labeled imager strand. (b) Super-resolution and diffraction-limited images of the microtubule network of fixed U2OS cells obtained with the smartphone-based microscope. The super-resolved image represents the localization density while the diffraction-limited image is the actual fluorescence intensity. Scale bar: 2 µm. (c) and (d) Comparison of super-resolved images of the microtubule network obtained with the smartphone-based setup (c) - magnified image of the region marked in (b) - and with the high-end microscope (d). Scale bar: 2 µm. (e) Line profiles across super-resolved microtubules as indicated in (c) and (d). Black solid lines: smartphone-based microscope. Pink dashed lines: high-end microscope.

## Discussion

We have demonstrated the feasibility of directly detecting single-molecule fluorescence using a simple, low-cost, portable microscope that leverages the image sensors and data handling capacity of mass-produced smartphones. We have validated this capability both in model DNA origami systems and in biological cells. This setup can be easily paired with any smartphone to detect single molecules, representing a significant leap towards universalizing quantitative sensing. We envision the smartphone-based microscope presented here will inspire new developments and biotechnological innovations, endowing single-molecule sensitivity to a wide range of applications such as digital bioassays, POC diagnosis, field expeditions, STEM outreach, and life science education.

Key to achieving direct detection of single-molecule emission was the design of the Total Internal Reflection (TIR) excitation and the use of raw RGB data. In contrast to previous works, we used a diode laser prism-coupled to generate high-angle TIR illumination. This configuration provides high irradiance while ensuring a short-range evanescent field (see Methods and Supplementary Information S8), effectively reducing the background signal. Analyzing raw RGB data allowed us to choose the most convenient detection channel, further enhancing the SBR. For example, the emission of single ATTO 542 molecules was detected with a 30% higher SBR when analyzing the green channel exclusively compared to the red and green channels combined (Figure 2e). The increase in SBR is even higher when compared to the total RGB signal. Analyzing the data in raw mode is also key for precise single-molecule localization.

Future iterations will focus on improving photon collection efficiency and measurement throughput. This could involve optimizing smartphone camera parameters and software or upgrading optical components at the expense of a small increase in the overall cost. For example, the photon collection efficiency could be improved by upgrading the NA of the objective lens, e.g., using a standard PMMA acrylic Fresnel lens. Using higher-quality interference optical filters in the detection instead of a single, colored glass filter would lead to a higher SBR.

With an estimated cost below 350 €, our results bring single-molecule measurements and assays within reach of any research laboratory, biotech company, or higher education institution. The setup requires minimal experience in optics, and a brief tutorial and short training make the microscope accessible to any interested user. Notably, the time-demanding super-resolution imaging measurements yielded comparable results when performed on a standard office desk and on an optical table, demonstrating the portable setup’s versatility (see Supplementary Information S9 for an exemplary super-resolved image).

Finally, our work paves the way for innovation in personalized assays with single-molecule sensitivity. The low cost and wide availability provide biotechnology enterprises, especially start-ups, with a new dimension to project their applications leveraging single-molecule sensitivity. Digital assays based on counting single molecules are easier to validate and calibrate reliably. Massively distributed assays based on smartphones can communicate with remote servers to validate and analyze data, harnessing the power of big data and AI tools. This could elevate diagnostics and disease prevention to unprecedented levels of efficiency.

## Methods

### Materials

Acetone: 133-VL54TE, Thommen-Furler. Isopropanol: 172-VL54TE, Thommen-Furler. Quartz coverslips: W9QA0490490170NNNNX1, Microchemicals. Photoresist and developer: AZ 1512 HS and AZ Developer from Microchemicals, respectively. Glass slides: 1000000, Paul Marienfeld. KOH pellets: 1310-58-3, Sigma-Aldrich. 50x TAE: J63931.K3, ThermoFisher. 1 M MgCl_2_: J61014.AK, ThermoFisher. 2-component removable glue: Elite Double 16 Fast, Zhermack. NeutrAvidin: 10443985, Fisher Scientific. BSA-biotin: A8549, Sigma-Aldrich. 10x PBS: J75889.K2, ThermoFisher. Double-sided adhesive tape: OCA8146-3, Thorlabs. Fiducial nanoparticles: 60 nm diameter gold nanoparticles, EM. GC60, BBI Solutions. 3,4-Dihydroxybenzoic acid (PCA): 37580-25G-F, Sigma-Aldrich, Protocatechuate 3,4-Dioxygenase from Pseudomonas sp. (PCD): 9029-47-4 P8279-25UN, Sigma-Aldrich. Cyclooctatetraene (COT): 138924-1G, Sigma-Aldrich. Trolox: 238813-1G, Sigma-Aldrich. Dimethyl sulfoxide (DMSO): 276855-100ML, Sigma-Aldrich. Agarose: 84004, Biozym Scientific GmbH.

### Buffers preparation

TAE12 buffer was prepared using 1×TAE and 12 mM MgCl_2_. 1×PPC imaging buffer was prepared using a PCA/PCD oxygen scavenging system and COT as a triplet quencher. In the case of the PPT imaging buffer, we used PCA/PCD plus Trolox as a triplet quencher. PCA aliquots were prepared: 154 mg of PCA in 10 ml of Mili-Q water, and pH was adjusted to 9.0 using NaOH. PCD aliquots were prepared: 8.3 mg of PCD (full bottle) and 11.3 ml of the following buffer (50% glycerol, 50 mM KCl, 1 mM EDTA, and 100 mM Tris). Trolox aliquots were prepared: 100 mg of Trolox, 430 μl of methanol, and 400 μl of NaOH 10 M in 3.2 ml of Milli-Q water. COT aliquots were prepared: 42 mg of COT in 2 ml of DMSO, further diluted in 6 ml of Milli-Q water. All aliquots were stored at -20 °C for up to 6 months. Imaging buffers were prepared using those aliquots and TAE12 buffer to reach final concentrations of 6 mM PCA, 20 nM PCD, and either 2 mM COT for PPC or 2 mM Trolox for PPT.

### DNA origami folding

We used single-stranded DNA M13mp18 from Bayou Biolabs as a scaffold. See Supplementary Information S11 and S12 for further details on the staples. The DNA origami nanostructures were folded with a 10-fold excess of staples to the scaffold (except for the oligos with fluorophores, which were included with a 1000-fold excess) in a folding buffer, TAE12, with a 20 h non-linear temperature ramp (ramp up to 95 °C, then go to 75 °C 5 min, to 65 °C 20 min, and ramp down 1 °C each 20 min until 20 °C) in a peqSTAR 2X, VWR thermal cycler. Gel electrophoresis was used for purification. The gel contains 0.8% agarose and TAE12 buffer. After adding the gel loading buffer (50% glycerol, 50% TAE12) to the DNA origami solution, the cooled gel ran for 3 h at 70 V. Afterwards, the gel was cut, and the DNA origami solution was extracted via squeezing. The DNA origami concentration was measured with a NanoDrop One^c^ Spectrophotometer, ThermoFisher Scientific.

### DNA origami sample preparation

Quartz coverslips were cut into 49 mm × 16.3 mm substrates with a laser cutter (Q400, Trotec) and left in acetone for 1 h. Next, they were rinsed with isopropanol and soaked for 1 h in fresh isopropanol. Then, the substrates were rinsed with Milli-Q water and dried with compressed nitrogen plus 5 min on a hot plate at 100 °C for water desorption. For wide-field/smartphone microscope correlation, 12 nm-thick chromium grids were created over the quartz substrate using photolithography (Direct Writing Laser system: μMLA, Heidelberg Instruments), sputtering (MiniLab 080, Moorfield Nanotechnology), and lift-off.

To remove any resist residue, quartz substrates with grids were cleaned by sonication in acetone for 10 min, followed by sonication in isopropanol for 10 min. Next, the coverslips were rinsed with Milli-Q water and dried with compressed nitrogen (purity 9N5). Then, substrates were exposed to ozone and UV radiation (PSD Pro Serie, Digital UV Ozone System, Novascan) for 10 min. Further, quartz coverslips were placed in a 3 M KOH bath for 10 min. Milli-Q water was used to rinse the quartz substrates, which were then dried with compressed nitrogen. Finally, substrates were exposed to UV/ozone to make the surface hydrophilic. To make the sandwich chambers, microscope soda lime glass slides were exposed to ozone and UV radiation for 10 min. Then, glass slides were placed in a 3 M KOH bath for 10 min. Milli-Q water was used to rinse the glasses, which were dried with compressed nitrogen. Finally, substrates were exposed to UV/ozone to make the surface hydrophilic.

Once the microscope slides and quartz substrates had been cleaned, 2 stripes of 4 layers of double-sided adhesive tape were applied to the quartz coverslip to form a channel of ∼300 μm height. Before closing the chamber, the quartz surface was functionalized with BSA-biotin/NeutrAvidin. First, the surface was incubated for 30 min using 100 μl of 0.5 mg/ml biotinylated BSA diluted in 1xPBS. Second, the surface was rinsed with 200 μl 1×PBS three times, followed by incubation with 100 μl of 0.5 mg/ml NeutrAvidin diluted in 1×PBS:Milli-Q (ratio 19:1) for 30 min. Next, the surface was rinsed three times with 1×PBS and one last time with TAE12. Next, a droplet of DNA origami solution was incubated to reach the desired surface density with the aid of the fixed dyes incorporated into the DNA origami design (see Supplementary Information S11). Typically, a 100 μl solution with 10 pM origami concentration in TAE12 was incubated between 10 to 30 min. Later, the sample was rinsed three times with 200 μl of TAE12. For storage, samples were left in TAE12 buffer, while PPT or PPC buffer was used for imaging. In the case of super-resolution experiments, fiducial nanoparticles were incubated after DNA origami immobilization, and dye-labeled oligos for DNA-PAINT were added to the imaging buffer. Finally, the chamber was closed with a cleaned glass slide using a 2-component removable glue. Both ends of the channel are glued to prevent oxygen exchange with the environment and buffer evaporation or contamination.

### U2OS cells sample preparation

A human bone osteosarcoma epithelial (U2OS) cell culture with immunolabeled microtubules was purchased from Massive Photonics. The cells were immobilized on a glass coverslip and sealed with an ibidi μ-slide I Luer 0.6 mm height single-channel slide. Immunostaining was done with a primary rat monoclonal antibody against α-tubulin and a secondary monoclonal anti-rat antibody conjugated with docking strands for DNA-PAINT. 90 nm gold nanoparticles were deposited on the surface as fiducial markers for drift correction. In the smartphone-based microscope, the sample was imaged with an imager concentration of 0.5 nM in a 1.5×PPC imaging buffer, in which TAE12 was replaced with 1×PBS. In the high-end microscope, we used an imager concentration of 10 nM in 1×PBS. For storage, the sample was rinsed 3 times with 200 μl of 1×PBS and left at 4 °C.

### Smartphone-based microscope measurements

Samsung Galaxy S22 Ultra, iPhone 14 Pro, and Huawei P20 Pro smartphones were used to detect a single molecule using our smartphone-based microscope. In the case of the Samsung smartphone, the telephoto camera was used; for the iPhone 14 Pro, the telephoto camera was used; and for the Huawei smartphone, the wide color camera was used. A low-cost colored-glass long-pass filter (10CGA-550, Newport) was used after the inexpensive 0.2 NA, 1.7 mm focal length, fish-eye objective lens (B07FXVJVDP, Richer-R) to collect the fluorescence signal and eliminate the residual scattered laser beam. The prism is a 4 mm diameter N-BK7 half-ball lens (#45-934, Edmund Optics) fixed with optical glue (NOA65, Thorlabs) under a cut microscope slide. Immersion oil (Type F, Leica Microsystems) was placed between the prism and the sample. The laser beam enters the half-ball lens at an incidence angle close to 80° to ensure the entire beam is totally reflected by the coverslip (see Supplementary Information S8). This angle also compensates for the refraction induced by the prism, as the whole laser beam cannot arrive perfectly perpendicular to the surface. The laser module (CW532-020F, Roithner LaserTechnik) has a wavelength of 532 nm and a power of 20 mW. The beam can be focused using the focusing lens incorporated into the laser module. In this work, the laser was focused on the sample plane, rendering an elliptical Gaussian beam with waists *ω*_*x*_ = 45.6 μm and *ω*_*y*_ = 27.6 μm (1/e^2^ of the peak intensity). Power on the sample plane was measured (S170C with PM100D, Thorlabs) to be (17.0 ± 0.5) mW, which is equivalent to a top-hat irradiance of 0.76 kW/cm^2^ in an area of 20 × 20 um^2^.

### High-end widefield fluorescence microscope

A schematic of the high-end custom-built wide-field fluorescence microscope is provided in Supplementary Information S10). It is based on an inverted Olympus IX83 body. Lasers with wavelengths of 532 nm and 640 nm (gem 532 and gem 640, Laser Quantum) were used to excite the samples. Each laser line was filtered with a clean-up filter (ZET532/10x, Chroma and ZET642/20x, Chroma) and spatially filtered with a 1x telescope to create Gaussian illumination. The two beams are aligned in a common path with a long-pass filter (RT532rdc, Chroma). Circularly polarized excitation is achieved using a polarizer (LPVISC100-MP2, Thorlabs) and a quarter waveplate (AQWP05M-600, Thorlabs). A neutral-density filter wheel was used to adjust the intensity. The two-color beam was enlarged with a 3.3x beam expander (AC254-030-A-ML and AC508-100-A-ML, Thorlabs), and then focused onto the back focal plane of the objective by a focusing lens (ACT508-300-A-ML, Thorlabs). This system is mounted on a motorized stage (LTM60-50-HSM, OWIS) that can be used to select wide-field, HILO, or TIR illumination conditions. A double-band dichroic mirror (ZT532/640rpc-UF2, Chroma) is used to direct the excitation light to the objective (UPLAPO100xOHR, 1.5 NA, Olympus) and let fluorescence light through. A double-band emission filter (ZET532/640m-TRF, Chroma) is used before the CMOS camera (C14440 ORCA-Fusion, Hamamatsu). The size of the detection PSF estimated as the width of the intensity profile at 1/e^2^ of its peak was *ω* = 2*σ* = (0.22 ± 0.03) μm. For single-molecule detection of the ATTO 647N, we used epi-illumination with a 640 nm Gaussian beam profile with a width (1/e^2^ of its peak was) of *ω* = (12.6 ± 0.1) μm, while for DNA-PAINT experiments we used a 532 nm Gaussian beam profile with a width of *ω* = (17.3 ± 0.1) μm in a TIR configuration.

### Recording and processing of videos and pictures with the smartphone

Videos and photos were recorded with MotionCam Pro: RAW Video app version 2.0.2-pro (available at Google Play) when using a Samsung Galaxy S22 Ultra smartphone. In the case of the iPhone 14 Pro, we used the Camera Pro by Moment app version 5.2.5 (available at the Apple store). The FreeDCam app version 4.3.22 was employed for the Huawei P20 Pro (available at GitHub). Videos acquired with MotionCam were recorded in MCRAW format and exported as a sequence of color (RGB) uncompressed DNG images. Videos acquired with FreeDCam were only recorded in RGB MP4 format. Both applications can acquire pictures directly in DNG. MP4 videos were converted to TIFF using Adobe Photoshop while DNG sequences were converted to TIFF using a custom-made MATLAB code. RGB videos were then split into red, green, and blue channels. Pixel outliers were first removed with Fiji (ImageJ2) software^33^.

The signal-to-background ratio (SBR), or Weber contrast, was calculated from one image as SBR = (*S* − *B*)/*B* where *S* and *B* represent the fluorescence and background intensities, respectively. First, the baseline of the smartphone camera was determined from a dark image and subtracted from all pixels. Next, to virtually increase the exposure time, we added the first 5 frames up to get an equivalent exposure time of 1.25 s. Then, for each detected molecule, we defined an inner region of interest (ROI) of 6 × 6 px^2^ and an outer ROI of 8 × 8 px^2^ centered around the diffraction-limited fluorescence spot. We obtained *S* after averaging the pixel intensities over the inner ROI. In like manner, *B* was obtained from the average intensity of surrounding pixels. The difference between the outer and inner ROIs produces a 2-pixel width squared ring from which *B* was calculated. Spots that showed outer ROIs overlapping other spots were not considered.

The signal-to-noise ratio (SNR) was calculated from single-molecule intensity traces as SNR = (*S* − *B*)/*σ*_*S*_, where *S* and *σ*_*S*_ are the time average and standard deviation of the intensity trace before the molecule photobleaches, and *B* is the time average of the background signal near the molecule. Traces were obtained from recorded videos after averaging the intensity of a 6-pixel diameter circular ROI around the diffraction-limited spots.

## DNA-PAINT imaging

We performed DNA-PAINT experiments using modified oligonucleotides that present a Cy3b fluorophore on one end and a fluorescence quencher on the other. The sequence of the imager strand and its complementary docking strand are 5’-BHQ2-AAGTTGTAATGAAGA-Cy3b-3’ and 5’-TTATCTCCTATACAACTTCC-3’, respectively. When hybridized with the docking strand, the 15 nucleotide-long imager strand has a pre-designed mismatch of 3 base pairs, which causes an average temporary bind of (1.05 ± 0.04) s, as shown in Supplementary Information S6. The imager concentration was 10 nM in 1×PPC imaging buffer for DNA origami imaging. We used an imager concentration of 0.5 nM in 1.5×PPC for the biological sample. Picasso software^29^ was used to analyze DNA-PAINT videos. Data was fitted using Maximum Likelihood Estimation and further filtered to keep only localizations with a localization precision below 31 nm. For rendering, localizations were un-drifted using nanoparticles as well as origami nanostructures.

## Data availability

Presented data is available at the Zenodo repository: https://doi.org/10.5281/zenodo.10979175. Further data is available from the corresponding authors upon reasonable request.

## Code availability

The code for DNG to TIF conversion with RGB channel extraction, implemented in MATLAB, and the custom Python-based software used to extract SBR from an image and SNR from videos are available upon reasonable request.

## Acknowledgements

M.L. and G.P.A thank A. Antonini from SmartMicroOptics for his helpful advice and fruitful discussions. G.P.A. and F.D.S. would like to thank Alan Szalai, Sebastián Giusti and Lucía Lopez for their support with preliminary samples. M.L. acknowledges funding from the University of Fribourg (Fonds de Recherche du Centenaire program). F.D.S. acknowledges the support from Agencia I+D+i, project PICT-2021-01216. G.P.A. acknowledges support from the Swiss National Science Foundation (200021_184687) and the National Center of Competence in Research Bio-Inspired Materials NCCR (51NF40_182881).

## Author Contributions

G.P.A. conceived the idea of the project. M.L. and N.F. designed and fabricated the smartphone-based setup. M.L., M.B., F.D.S and G.P.A. designed the experiments. M.L. and M.B. did the experiments. S.K., K.K. and M.L. designed, synthesized, and characterized the DNA origami nanostructures. M.L. and M.B. analyzed the data and made the figures. M.B., F.D.S. and G.P.A. coordinated the project. M.B., M.L., F.D.S. and G.P.A. wrote the manuscript with input from all the authors.

