## Supplementary Information for "Direct single-molecule detection and super-resolution imaging with a low-cost portable smartphone-based microscope"

Corresponding authors:

#### S1. Smartphone-based setup configurations

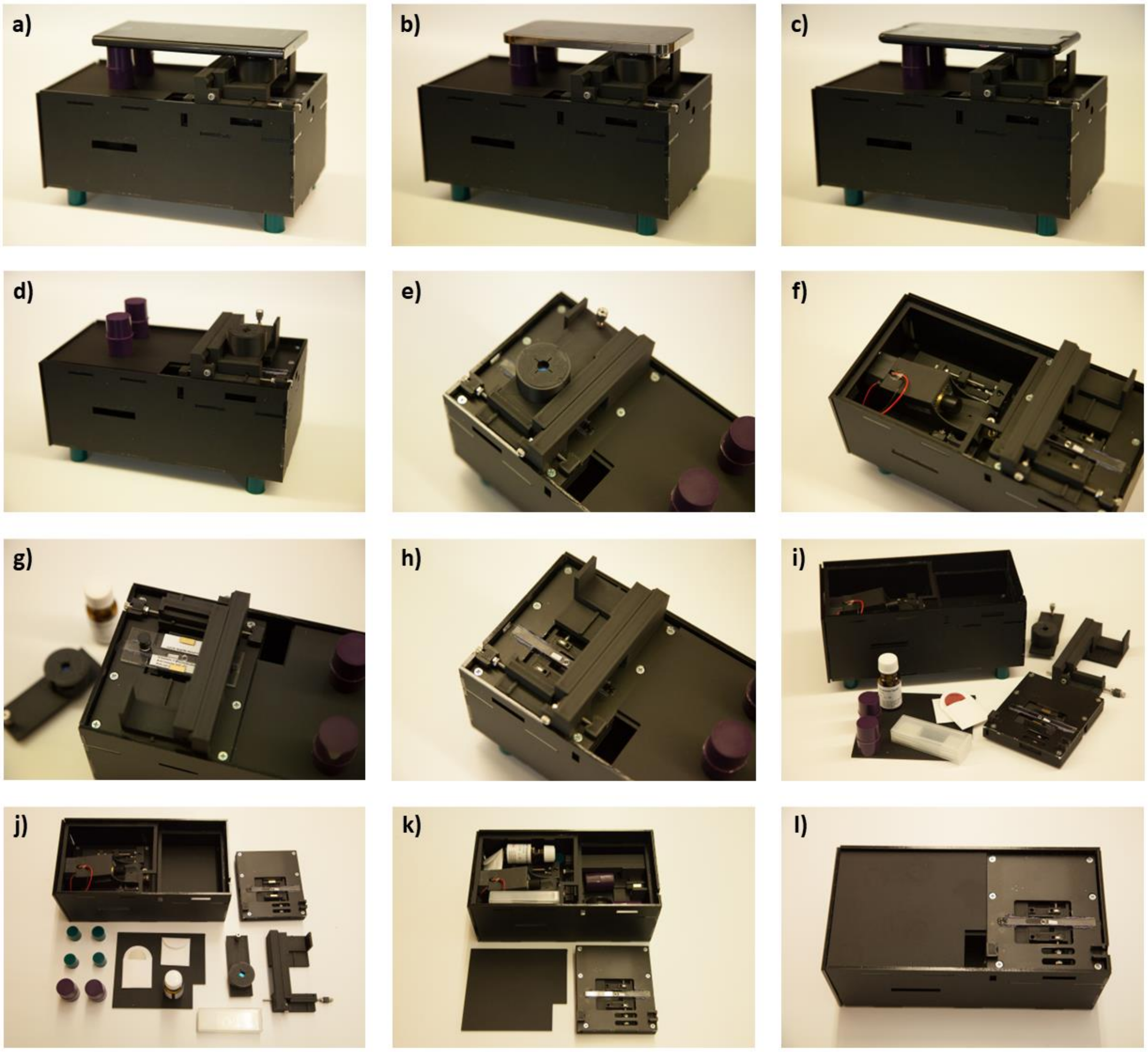

*Figure S1: Different configurations of the Smartphone-based setup featuring (a) Samsung Galaxy S22 Ultra, (b) iPhone 14 Pro, (c) Huawei P20 Pro, and (d) no smartphone. (e) Top view of the objective stage. (f) Half-cover and objective removed. The laser stage and sample stage are visible. (g) Mounting a sample. Magnets (metallic yellow rectangles) hold the sample to the sample stage. (h) Top view of the sample stage. (i) and (j) Smartphone-based setup components, immersion oil, filters, sample box, and slip-resistant feet. (k) and (l) Smartphone-based setup ready for transportation.*

#### S2. Smartphone-based setup price list

*Table S1: Price list of relevant components.*

| Component | Price (€) |
|---|---|
| Objective (B07FXVJVDP, Richer-R) | 11.00 |
| Half-ball lens (#45-934, Edmund) | 45.00 |
| Emission filter (10CGA-550, Newport) | 57.75 |
| Optical glue, a fraction (NOA65, Thorlabs) | 2.00 |
| Glass slides (a few of them) | 1.00 |
| Laser + heatsink (CW532F-020F, Roithner LaserTechnik) | 70.35 |
| Electronics (connectors, cables, potentiometer, switch, fan, LED, voltage regulator) | 26.30 |
| Battery (EB-P3300, Samsung) | 53.55 |
| Silicone feet (Ecoflex 00-50, Smooth-on) | 4.83 |
| Frame (3D-printed parts, laser-cut parts, glue) | 15.79 |
| Miscellaneous (screws, knobs, bolts, threaded insert, magnets) | 33.76 |
| **Total** | 321.33 |

#### S3. TEM of the 2LS and 8HB DNA origami nanostructures

For TEM imaging, 5 µL of sample solution were dropped onto formvar-coated grids (300 mesh Cu, 22-1MHC30-50, Micro to Nano). After 1 min, the solution was removed with a paper filter, and the sample was negatively stained for 10 s using 2% Uranyl acetate solution. TEM images were acquired with a FEI Tecnai Spirit microscope with an accelerating voltage of 120 kV.

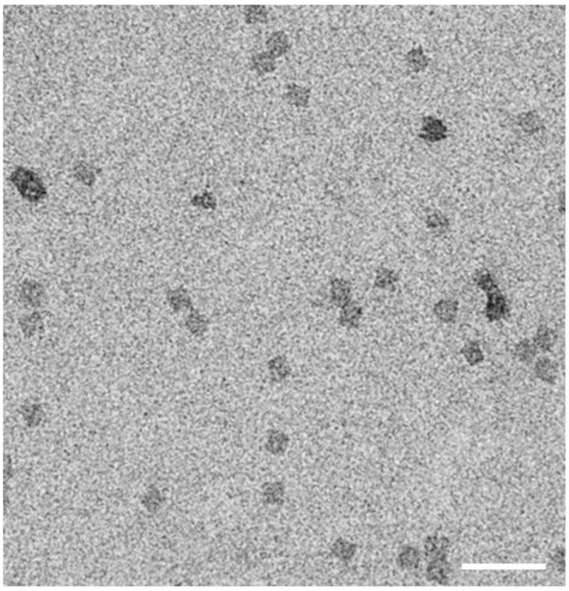

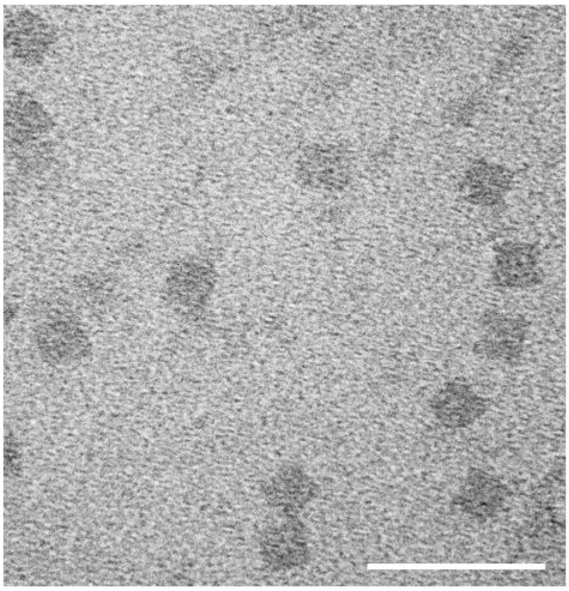

*Figure S2: TEM images of purified 2LS nanostructures. Scale bars: 200 nm.*

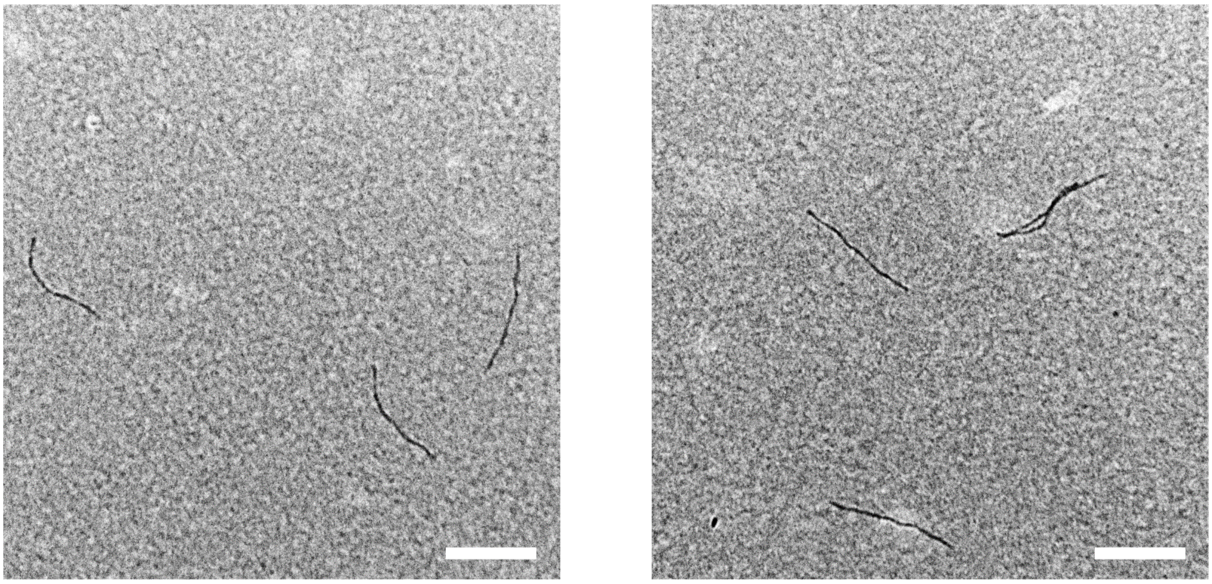

*Figure S3: TEM images of purified 8HB nanostructures. Scale bars: 200 nm.*

#### S4. Characterization of dye incorporation in the 2LS DNA origami nanostructures

To characterize the percentage of 2LS origami nanostructures containing both dyes (ATTO 542 and ATTO 647N) and those having only one, the samples were studied using the high-end microscope and 1×PPC imaging buffer. First, the nanostructures were excited with a 640 nm laser until the red fluorophores were photobleached, and, subsequentially, the same region of interest was excited with a 532 nm laser until the green ones were photobleached too. Intensity traces of fluorescent spots were analyzed. Traces showing only single steps were considered. All other cases were discarded, e.g., two steps, blinking, or spots that didn't photobleach. After counting single molecules that spatially colocalize, we found that 87% of the nanostructures had single ATTO 542 molecules (Figure S4a), 85% had single ATTO 647N molecules (Figure S4b), and 72% had incorporated both dyes (Figure S4c). This result agrees with the saturated incorporation efficiency reported by Strauss *et al.* for the maximum explored staple excess over the scaffold[1]. In Figure S4d, colocalization between the smartphone-based and high-end microscopes is shown. Due to the PSF size of the smartphone-based setup, 1:1 and 2:1 cases were taken into account. The data correspond to 41 fluorescent events registered in Figure 2f of the main text, out of which 87% was considered to come from DNA origami nanostructures.

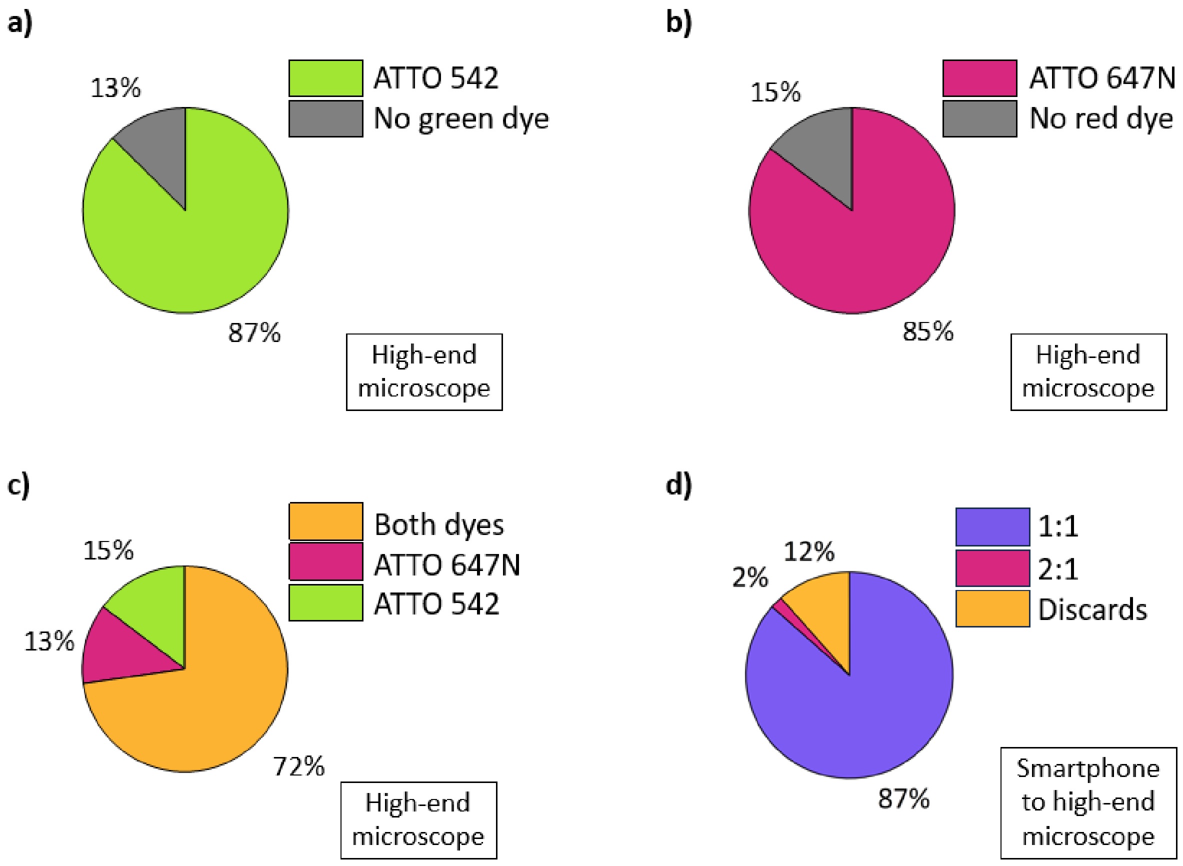

*Figure S4: Single-molecule incorporation in 2LS using the high-end microscope. Percentage of nanostructures showing a single ATTO 542 (a), a single ATTO 647N (b), and both molecules (c). A total of 143 nanostructures were analyzed, of which 104 presented both dyes, 122 presented single ATTO 647N molecules, and 125 presented single ATTO 542 molecules. (d) Colocalization between microscopes. Legend N:M represents the number of ATTO 542 photobleaching steps observed in the*

*smartphone-based setup (N) against the number of ATTO 647N photobleaching steps observed in the high-end microscope (M).*

#### S5. Direct single-molecule detection with more smartphones

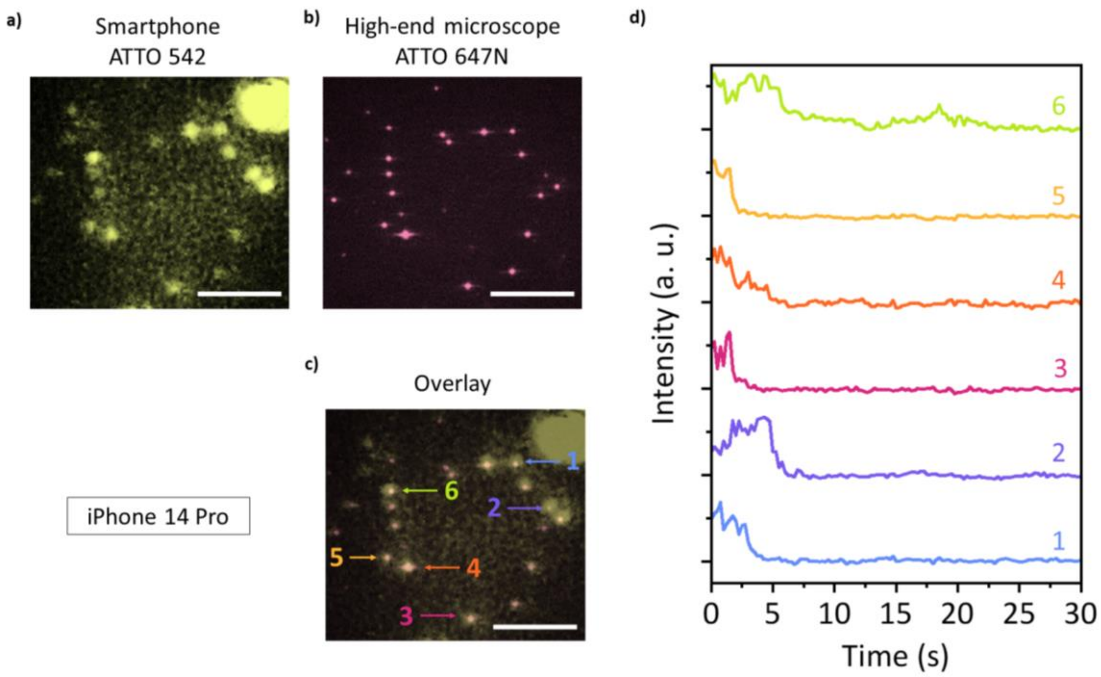

*Figure S5: Direct single molecule detection with the smartphone-based microscope using an iPhone 14 Pro. (a) Fluorescence image of the 2LS origami on quartz, detecting only the ATTO 542 with the smartphone-based microscope and (b) the ATTO 647N with the high-end microscope. (c) Overlay of images (a) and (b). Arrows and numbers indicate intensity traces plotted in (d). (d) ATTO 542 intensity traces vs. time. They correspond with the 2LS nanostructures indicated in the overlayed images shown in (c). Scale bars: 10 µm.*

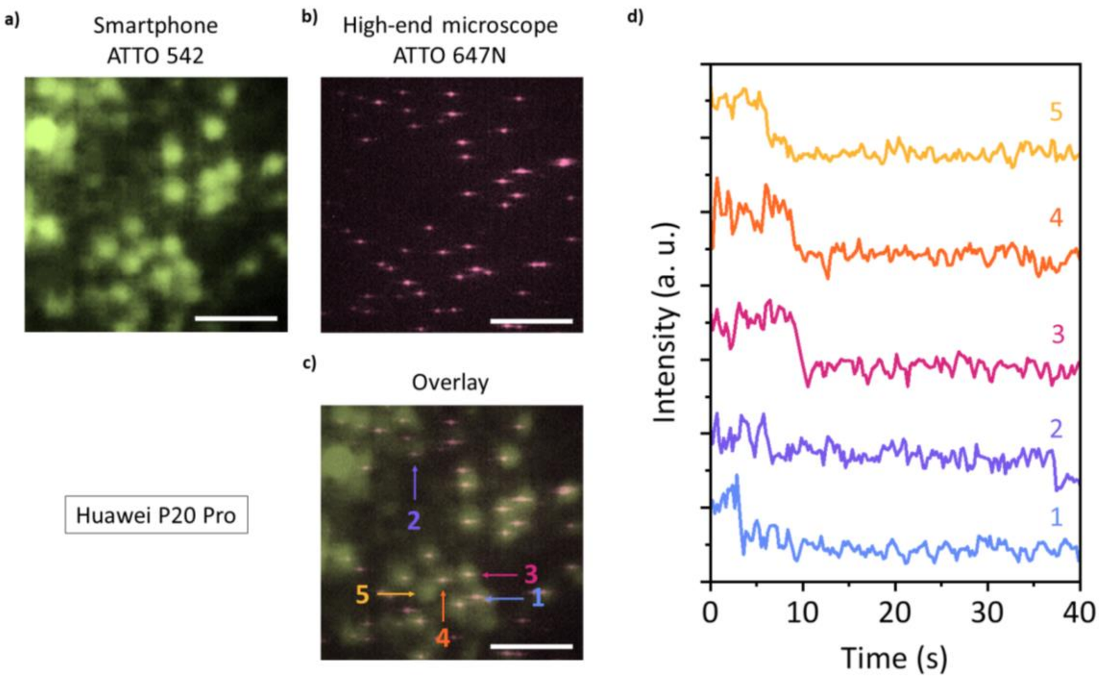

*Figure S6: Direct single molecule detection with the smartphone-based microscope using a Huawei P20 Pro. (a) Fluorescence image of the 2LS origami on quartz, detecting only the ATTO 542 with the smartphone-based microscope and (b) the ATTO 647N with the high-end microscope. (c) Overlay of images (a) and (b). Arrows and numbers indicate intensity traces plotted in (d). (d) ATTO 542 intensity traces vs. time. They correspond with the 2LS nanostructures indicated in the overlayed images shown in (c). Scale bars: 10 µm.*

#### S6. Kinetics and optimization of DNA-PAINT experiments

Binding events follow a mono-exponential distribution (Figure S7) at fixed room temperature (21 °C). Data was fitted assuming a mono exponential probability density function $pdf(t) = \left(\frac{1}{\tau}\right) e^{-t/\tau}$ in a Maximum Likelihood Estimation problem. We found an estimated mean binding time $\tau$ of (1.05 ± 0.04) s. A custom-made Python-based code was used (available at https://github.com/marianobarella/DNA-PAINT_thermometry).

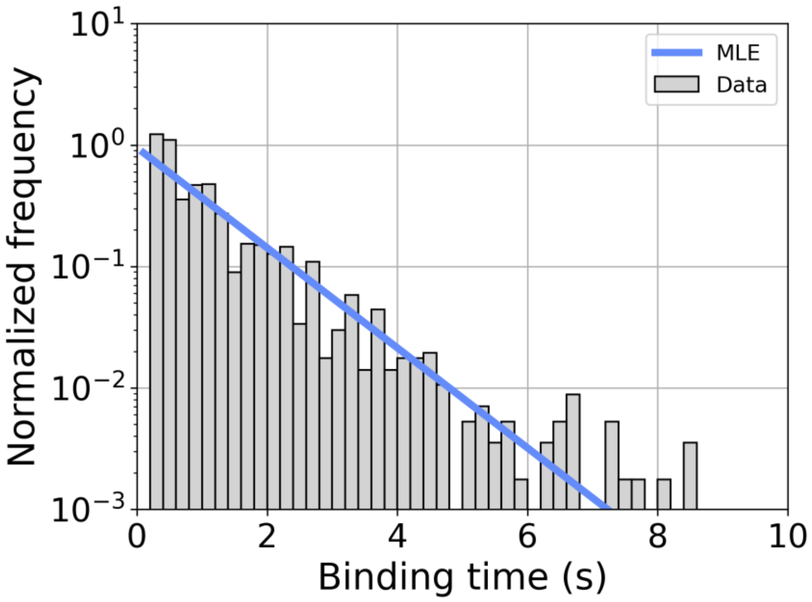

*Figure S7: Normalized frequency of 2836 DNA-PAINT binding events of the different binding times. The blue line represents a mono exponential probability density function with τ = 1.05 s obtained after applying the Maximum Likelihood Estimation method.*

Image quality was improved after averaging three frames. Figure S8a shows a comparison of ten fluorescent events - corresponding to Cy3b single molecules - with and without averaging. After applying Picasso's Localize[2] module to the original and the averaged video with the same parameters (identification, photon conversion, and fit settings), we found that the localizations' PSF sizes are more uniform (Figure S8b). When rendered, the drift-corrected localizations of the averaged video exhibit a more precise look than the original video, as shown in Figure S8c.

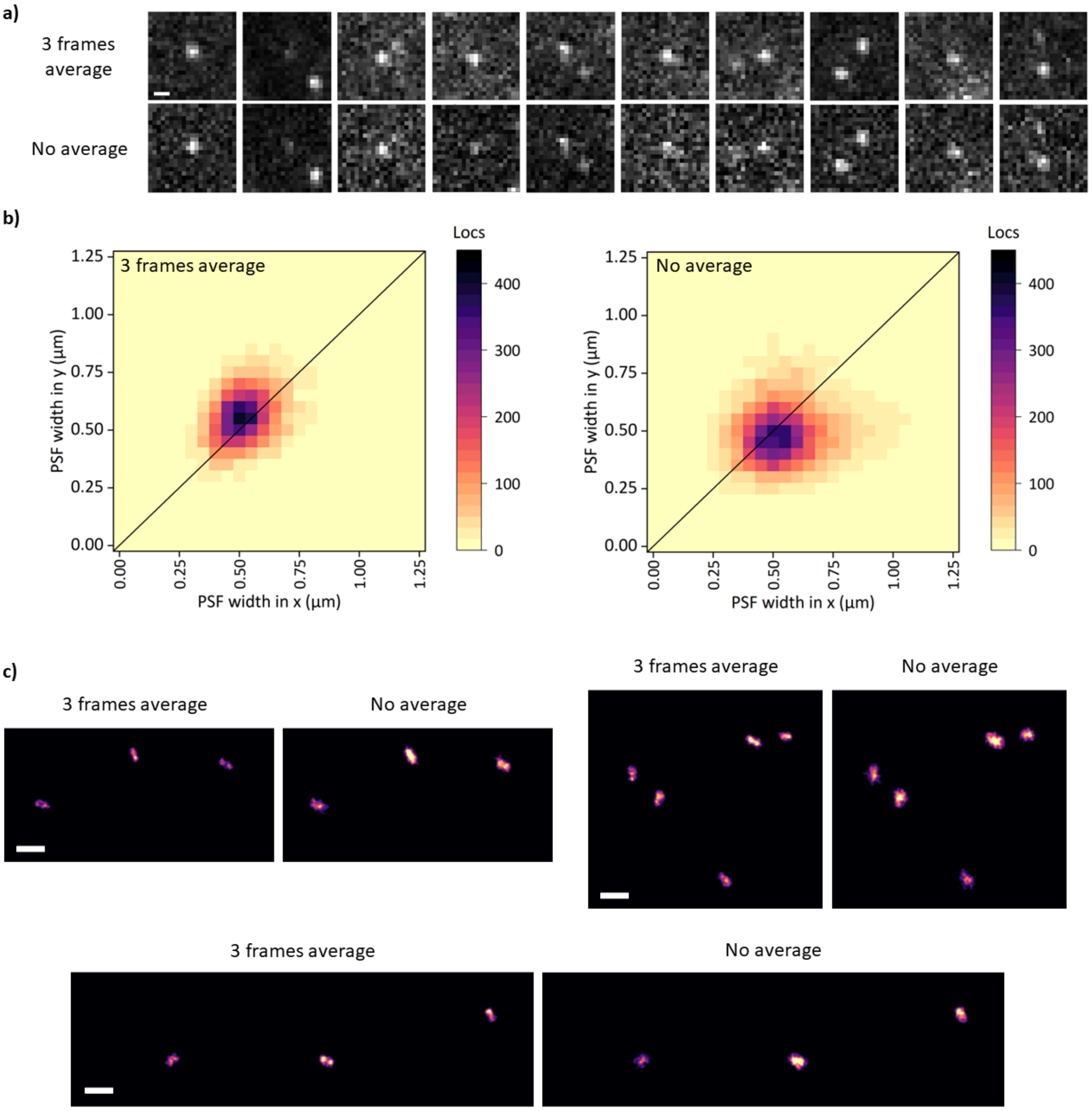

*Figure S8: Improving image quality by frame averaging. (a) One-to-one comparison of 10 single molecules. (b) PSF width distribution in x and y. The averaged video exhibits a lower dispersion, meaning the spots are more round and similar to each other. (c) Super-resolved images. The same analysis pipeline was applied to both three frames of averaged video and no averaged (original) video. Nanostructures are sharper when performing the average. Scale bar: 1 µm.*

To maximize the number of photons per frame, which in turn improves the localization precision, we tested Cy3b-labeled imagers with two modern oxygen scavenging and triplet quencher systems on the high-end microscope[3,4]: PCA-PCD/Trolox (PPT) and PCA-PCD/COT (PPC). See online Methods for details on their preparation. Figure S9 presents distributions of detected photons per frame, signal-to-background ratio (SBR), signal-to-noise ratio (SNR), and radial localization precision. PPC buffer presents a higher photon per frame and a higher localization precision[5]. SNR and SBR show no significant differences. PPC buffer was chosen for the DNA-PAINT experiments.

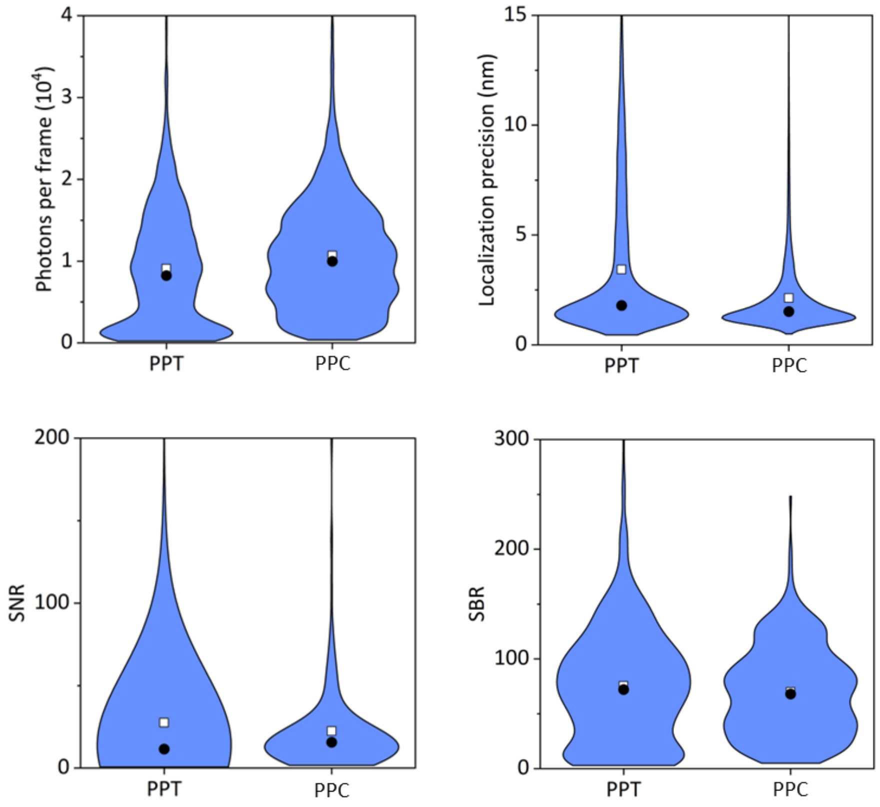

*Figure S9: Comparison between PPT and PPC imaging buffers. Distribution of photons per frame, SBR, SNR, and localization precision for 816 single molecules photostabilized with PPT and 556 with PPC. The average (white square) and median (black circle) are shown.*

### S7. Resolution estimation

To estimate the achieved resolution, we used a parameter-free estimation based on the decorrelation analysis presented by Descloux[6–8]. To be able to run the analysis over SMLM data, it is necessary to render the super-resolved image from the list of localizations. Selecting a pixel size (or even testing several values, as shown in Figure S10) is also essential. In this work, we chose the fixed Gaussian rendering method. That means each localization is represented by a Gaussian PSF centered on the localized coordinates with the same $\sigma$. We set $\sigma$ to the localization precision we found when doing DNA-PAINT, $\sigma$ = 86 nm for the smartphone-based setup and $\sigma$ = 24 nm for the high-end microscope (see Section *Super-resolution benchmark with DNA origami models* in the main text). We ran the analysis on the microtubule imaging data shown in Figure 4 (see main text). The number of localizations was $3.8 \times 10^4$ for the smartphone-based setup and $6 \times 10^6$ for the high-end microscope. The estimated resolution is approximately 210 nm for the smartphone-based setup and 66 nm for the high-end microscope.

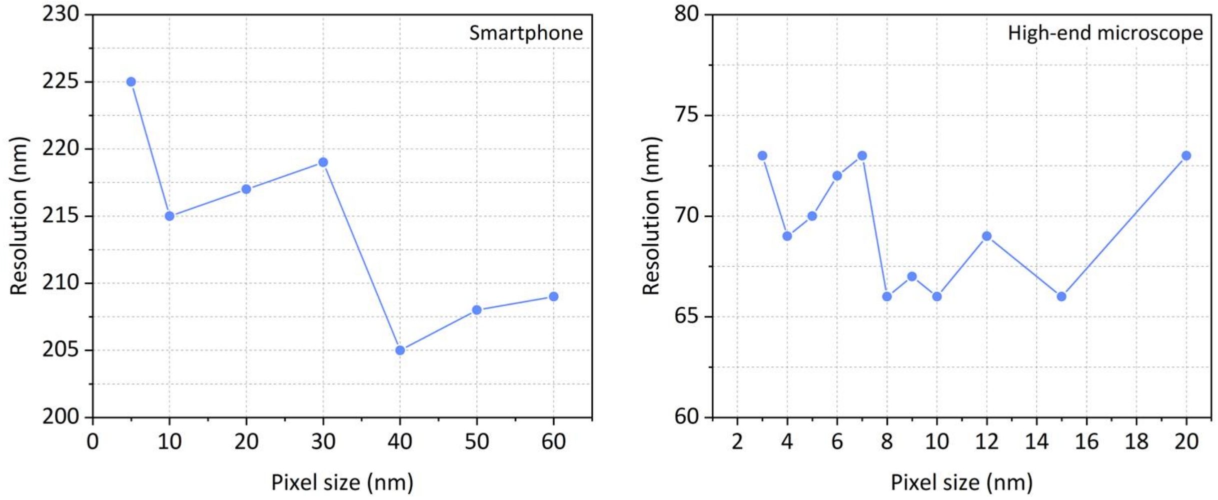

*Figure S10: Estimated resolution as a function of the pixel size for both setups.*

#### S8. TIR excitation design simulations

When focusing a collimated beam, total internal reflection is successfully achieved when all rays are effectively reflected, as shown in Figure S11. Geometrical optics raytracing simulations were performed using a 532 nm beam at an incident angle $\theta_i$ of 80°, a glass substrate ($n_i$= 1.52), a water medium ($n_i$= 1.33) and a critical angle $\theta_c$ = 61°. We used the Optics Workbench module (https://github.com/chbergmann/OpticsWorkbench) of the open-source software FreeCAD, version 0.20.2 (https://www.freecadweb.org/).

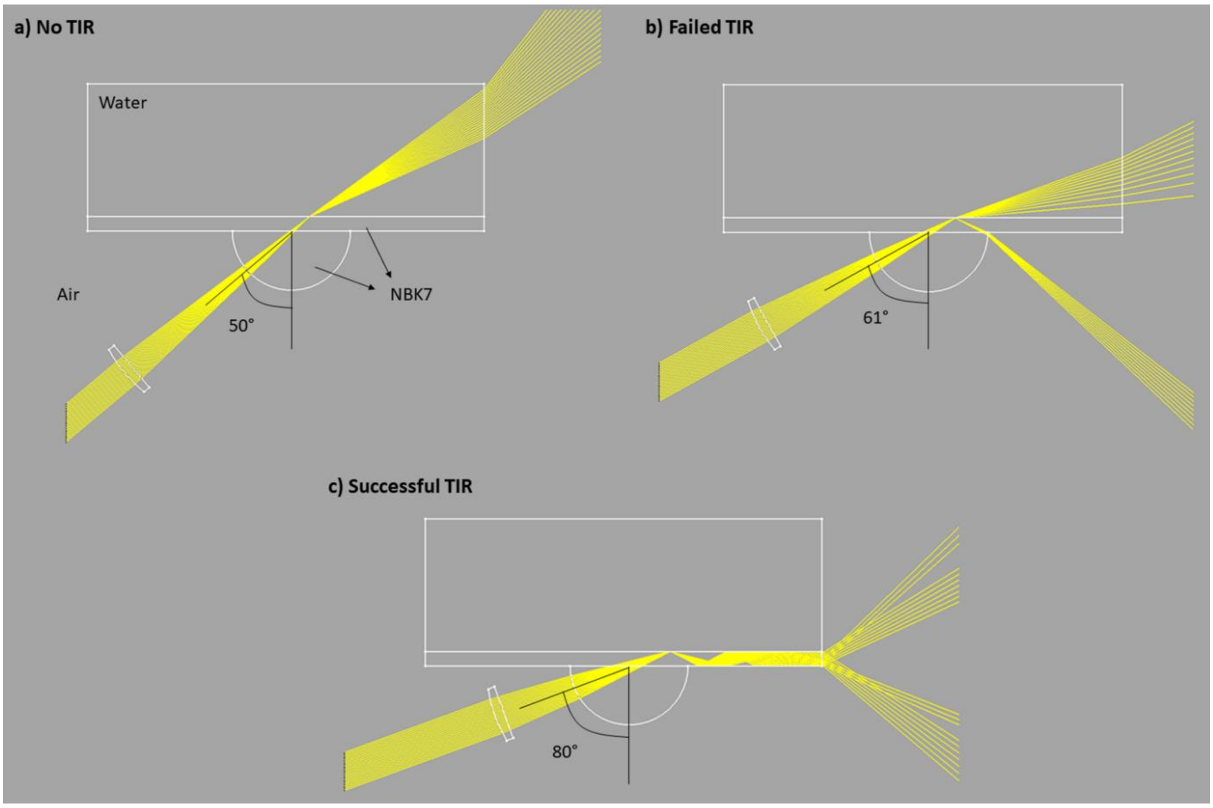

*Figure S11: Raytracing of the prism-type TIR illumination implemented in the smartphone-based setup. Different angle of incidence: (a) 50°, (b) 61° and (c) 80°.*

Finally, we can calculate the penetration depth of the evanescent field intensity as[9]:

$$d = \frac{\lambda}{4\pi} \frac{1}{\sqrt{n_i^2 sin^2(\theta_i) - n_s^2}}$$

For our smartphone-based microscope configuration, the penetration depth $d$ is 80 nm.

#### S9. Super-resolution on an office desk

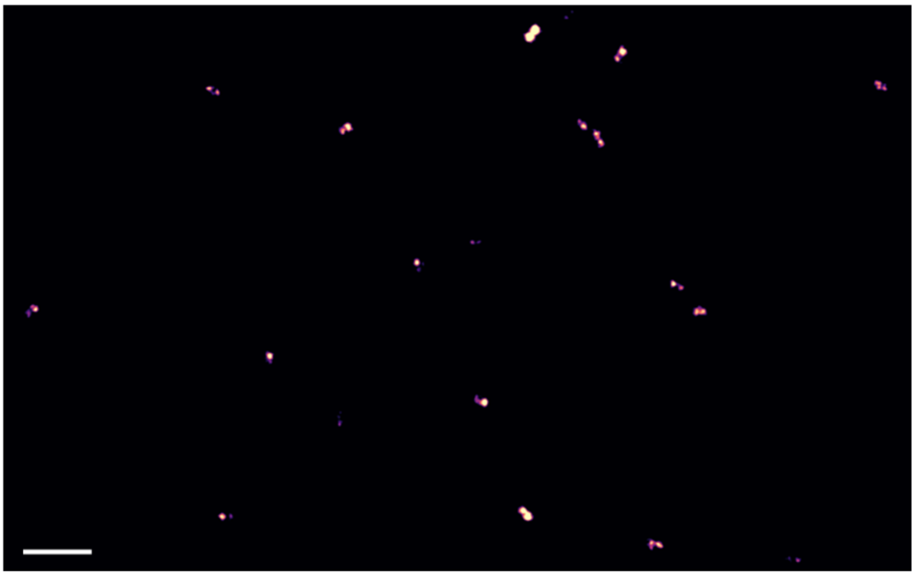

*Figure S12: DNA-PAINT supper-resolved localization image. Several 8HB nanostructures in aqueous solution. The measurement was performed on a standard office desk for ~3.5 h (see Supplementary Information S13). Scale bar: 2 µm.*

#### S10. High-end widefield fluorescence microscope

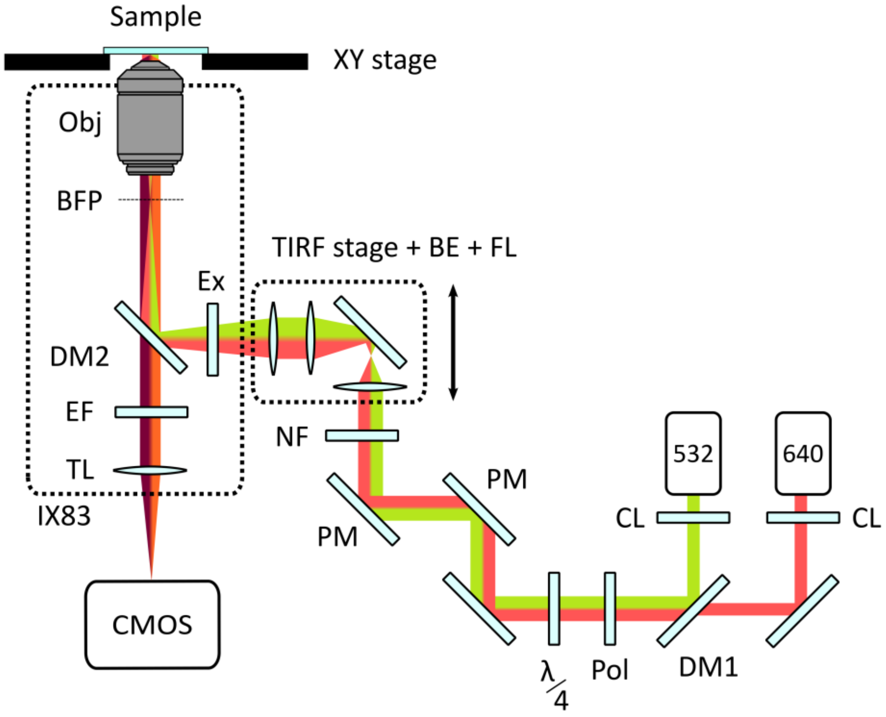

*Figure S13: Scheme of the High-end Widefield Fluorescence Microscope. CL: clean-up filter. DM1/2: dichroic mirror. Pol: polarizer. PM: periscope mirror. NF: neutral filter. BE: beam expander. FL: focusing lens. Ex: excitation filter. EF: emission filter. BFP: back focal plane. TL: tube lens.*

#### S11. 2LS DNA origami staples list

DNA origami nanostructures were designed with cadnanoSQ v0.2.4 (available at https://cadnano.org/legacy.html).

*Table S2: List of staples for the origami core.*

| Staple number | Sequence (5' to 3') | Nucleotide length |
|---|---|---|
| 1 | TCCTTTTGATAAGAGCATCAAGAAAACAAA | 30 |
| 2 | AAAACCAAAATAGCGGTGTGATAAATAAGG | 30 |
| 3 | GCTTAGAGCTTAATTGATTACCTGCTTTTTCA | 32 |
| 4 | GAAGTTTTGCCAGAGGTGACCTAATGGCTCAT | 32 |
| 5 | CCATCACCACGTGGCGGCGCTAGGGCACGTAT | 32 |
| 6 | CGCGTACTGGTAATATGAGTAAAAGACCTGAA | 32 |

| | | |
|---|---|---|
| 7 | AAAAAAGGACTTTCAAGCCTGTAGGCCACCAC | 32 |
| 8 | CCCCTAAGAGAATATAAAGTATTTTCGAGCCAGTAACCCC | 40 |
| 9 | CCCCTGTGAAATTGTTATCCGCGGGAAGGTGCAAGGCTTGACCGT | 45 |
| 10 | ATTACCATACGGAAATGTTTACCACATACATA | 32 |
| 11 | ATAATAACTAGCAATAGTCAGAGGAATGAAAA | 32 |
| 12 | AAACACTCGAAAGAGGCAGGGAGTATATATTC | 32 |
| 13 | AGAATAGATTTTTTCACATAACCGTAAAGGCC | 32 |
| 14 | AGGCGCATAACTAATGTTAAGAACATTTAATG | 32 |
| 15 | GATTATACGGATTAGAAATTTCATGGTTATAT | 32 |
| 16 | AGTCTCTGAATTTACCGAGCCGCCTCCTCAAG | 32 |
| 17 | CAAAGTTAAGAAAAGTGACGGGAGCATAAAAA | 32 |
| 18 | CGCCTGCATACATTTTACCCTTCTGAGTCTGT | 32 |
| 19 | CCCCTCAATAACCTGTTAACATTATGACCCTGTAATCCCC | 40 |
| 20 | CCCCAACAAAGCTGCTCATTCAGTAAAACGAACTAACCCC | 40 |
| 21 | CCCCCGGAACAACATTACTGCGGAATCGTCATAAATCCCC | 40 |
| 22 | CCCCTCTTTCCAGACGTTAGTAAATCAGCTTGCTTTCCCC | 40 |
| 23 | TGCCTGCACGACGGCCTTCTGGTGTCCAGCCA | 32 |
| 24 | GAGTGTTGTTCCAGTAATCGGAAGGGCAAC | 30 |
| 25 | AAGACTTCAAATATCGCGAGAAAAAGCAAAAG | 32 |
| 26 | CCCCATTCATTGAATCCCCCTCATTTACCCTGACTACCCC | 40 |
| 27 | CCCCAAGCGAAAACCGTCTATCAGGCCCC | 29 |
| 28 | AATTAATGTTATTTCAGTACCAAATAGCTATA | 32 |
| 29 | TAGCAAAAGAGAATCGGACAGTCAAATTCGCG | 32 |
| 30 | CATCAATTCTACTAATTACCAAGTATGCAATG | 32 |
| 31 | TTACTAAAAGGGAGCAATACTTCTTTGATTACCCC | 35 |
| 32 | GCCGGCGACAAATCAAACTCCAACGTCAAAGG | 32 |
| 33 | TAATGAGTGCGCTTAGAGAAGTGGCTTGCA | 30 |
| 34 | GACAAAAGATCGATACCGGAAAACCGGAAC | 30 |
| 35 | TCTGGCCTTGAGCGAGGATCGCACCCGGAAAC | 32 |
| 36 | CCCCATCAGAGCGGGAAGGGAAGACCCC | 28 |
| 37 | CCCCACTTTTGCGGGAGAAGCCTCCGGAGAGGGTAGCCCC | 40 |
| 38 | AATTTAGTCATTCCTTTACGAGGAGGTTT | 29 |
| 39 | AGATTAAACGCTCATTTAGTAAAGGTAA | 28 |
| 40 | CCCCAATCGGCTGTCTTTCCTTAGCAG | 27 |
| 41 | CCCCAAAATCCTGTTTGATGGTCCAGCTGCATTACCCC | 38 |
| 42 | CATGTTCATAGATAAGTCGAGAACTCATTACC | 32 |
| 43 | TACGTTAATGAATAAGCCGGATATACCTGCTC | 32 |
| 44 | AAACTAGCAAAATCTGGATTTAGTCAAAAG | 30 |
| 45 | AAAAGCGTAGATATTTTAAAAGTTTGAGTAACCCC | 35 |
| 46 | ATCAACAAGCTAATGCCATAGTAACGACGATA | 32 |
| 47 | ATTCACTTGCGGCATGTAGAAACCAATCAATCCCC | 35 |
| 48 | TAGCGACATTCATATGTATTCATTGAAACGCA | 32 |
| 49 | CCTAAAACATCTTTGACCGAACTGTACAGACC | 32 |
| 50 | GTAACAGTACCGCCAGGAATAGGGGATTTT | 30 |
| 51 | CCCCCCAAATAAGAAACGATTTTCGGT | 27 |
| 52 | AGACAGCCCGATAGTGGAGCCTTAAACGGG | 30 |
| 53 | CCCCTTTAGCGAACCTCCCGTAAGAACGCGAGGCGTCCCC | 40 |
| 54 | CGTTTTTAGACAGATCTTTAATCCCTGTTT | 30 |
| 55 | ACCACGGAACCGTAAACTAAAGGAGCCGAA | 30 |
| 56 | ATTAAATGTCCTGTAGTCATATGTTAATCGTA | 32 |
| 57 | CAAATAAAATATAAAATACCGAACGAACCACCCCC | 35 |
| 58 | ATAGAAAAGAATCAAGAATCACCACAGAACCG | 32 |
| 59 | TACATGGCCGCGTTTCACCCTCAGAGCCGCC | 31 |
| 60 | CTGTCGTGGGTTCCGATCCACGCTGGTTTGCCCCAGCAGGCGCCCC | 46 |
| 61 | TGCGCTCAAAAAGAATCCGCCTGGCCCTGAGA | 32 |

| | | |
|---|---|---|
| 62 | TTTTAGAAATATTCAATGCCTGAGCAGGAAGA | 32 |
| 63 | AAAGGAATTACGAGGAGAACGCGATTGTGA | 30 |
| 64 | AATTATCAAATATCAAAGATTAGACGCGAACT | 32 |
| 65 | CCCCCAGCAGAAGATAAAACAGATTAT | 27 |
| 66 | AGCTCAACTGGGGCGCGAGCTGACAGAGCAT | 31 |
| 67 | TCATAGCCCCCTTATTGCCACCCTGTAGCACC | 32 |
| 68 | CCTACCAATCGTCGCAGTACATAGACTAC | 29 |
| 69 | TCATAATCAAAATCACCGGACCCC | 24 |
| 70 | GAAATACCACAGTGCCCTTTAATGGCCGTCAA | 32 |
| 71 | CGCAAGACAAAGAACGCGTTTTAATGCGGATG | 32 |
| 72 | ACCCATGTATAAGTTACTTGAGCTTGCTCA | 30 |
| 73 | CCCCATTGAGGAAGGTTATCTCAACAGTTGAAAGGACCCC | 40 |
| 74 | GTGCATCTGCAAATATCAGCTCAGCTGATA | 30 |
| 75 | GCTAAACACTCCAAAATGCGCCGAGCAGCGAA | 32 |
| 76 | GCACCCAGAACGAGCGGAGAATAAAATTAACT | 32 |
| 77 | GTACCAGGTAGGATTAGATACAGGAAGCGTCA | 32 |
| 78 | AATCCTTTCAACTAATACCCTCAAATTAACAC | 32 |
| 79 | AATAATAAAAGGAACACACTGAGTCAATAGGA | 32 |
| 80 | TATCATCGATGAGGAAGAGGGTAGTTAAACAG | 32 |
| 81 | GCTTTTGCTAAGGGAACCCCCAGCGGGCTTGA | 32 |
| 82 | CGCGCAGATCATTTCACTGAATATAAGCCCGA | 32 |
| 83 | AGGCTATCATCGGTTACGCAAGGTTTGACCA | 31 |
| 84 | CCAACGCTCTACAATTCGTAGGAAAAGCAAGC | 32 |
| 85 | CCCCTTATTACGCAGTATGTTAGTATC | 27 |
| 86 | AGAAGGATCGGATAAGAGCAAGCCTTCGTCAC | 32 |
| 87 | CCCCCACAAACAAATAAATCCTCATTAAAGCAGGTCATCTGAAAC | 45 |
| 88 | CCCCGAGGACTAAAGACTTTTTCCCTGATAAATTGTCCCC | 40 |
| 89 | CTGAGGTCTGAGAAATCAATATATGTGAGTGCCCC | 35 |
| 90 | CCCCGTCGAAATCCGCGTCATTACCCAAATCAACGTCCCC | 40 |
| 91 | ATTATCACGCCAGCAATTTGCCTTATTTTCGG | 32 |
| 92 | CCACATTCAGGCTGGCAGTAAATTGATTATAC | 32 |
| 93 | GGCAATTCATCATACGTAAAGATAAGTATT | 30 |
| 94 | CCCCATTTATCAAAATCATAGAAGAGTCAATAGTGACCCC | 40 |
| 95 | AAATGGAGGTGAGGCGCCATTACTTTAGG | 29 |
| 96 | CCTGAGTAAGGCCGGAATGAACGGACCCCGGT | 32 |
| 97 | GAACACCCAACATATAAAACCGAGAAAGGTGA | 32 |
| 98 | AATCAGTGAGAATCCTATGCGCCGAACCACCA | 32 |
| 99 | CTTGATACCTCATAGTCTGTATGTGTATCA | 30 |
| 100 | ATCAGGTCAATGCTTTGTCCAATATTACAGGT | 32 |
| 101 | GAGTAATTTTAGTAATGACCATATCTGCGAA | 31 |
| 102 | AAGGTGGCTGAACAAAGCTATCTTGAGCCTAA | 32 |
| 103 | CTATTAAACGGTCACGGAGCTTGAACAGGAAC | 32 |
| 104 | GTATCGGCAGAAAAGCTCAAAAATAATCACCA | 32 |
| 105 | GAGAATCGTCCTTGAATTAGGTTGTTGAATTA | 32 |
| 106 | GAACCTCATCATATTCTCGACAACAACAGTAC | 32 |
| 107 | CCCTCAGAGCCCGTACCTATTATGACGATTGGCCTTGATATTCCCC | 46 |
| 108 | CCACCCTCAGAGCCACTCATCGGCTAGCGTCA | 32 |
| 109 | GCCTTGCTATGGTTGCGATTTTAGCGGGGAAA | 32 |
| 110 | GGAGGCCGCAAATTAAAGAACTCAATCGTCTG | 32 |
| 111 | CCCCGTCACACGACCAGTAAATTGGCAGATTCACCACCCC | 40 |
| 112 | AAGTGTAGGAACGTGGGTTTTTTGGGGTCGAG | 32 |
| 113 | AAAGCTAAAGGTCATCCGTTCTATTTTTTA | 30 |
| 114 | GTCAATCAGGGATCGTACTACGAAATCTCCAA | 32 |
| 115 | GCCAGCCAGCACCGCAGTGCCAATTTTTAT | 30 |
| 116 | TATACCAGGAACGAGTTGACCTTCCGCAGACG | 32 |

| | | |
|---|---|---|
| 117 | GGTACGCCAGGCCACCCCAGAACACGCTCATG | 32 |
| 118 | CGAATTATGGCGAGTAGATTTAGATAAAAAT | 31 |
| 119 | TAATATCCTGTCCAGAAACAACGCGCTTAATT | 32 |
| 120 | AGTAATTCCATCCTAAAAGAACGGTATAGAAG | 32 |
| 121 | TGGGTAACTCGAATTCCCACACAACGGGAAAC | 32 |
| 122 | CTATCATACGCACTCATCCTGAACGCTATTTT | 32 |
| 123 | AAACACCATCAGGACGTTGAGATTGTAAAATG | 32 |
| 124 | CCTTAGAATTCTGAATTATACAGTTCGTATTA | 32 |
| 125 | TTTTCATTATGTTTTTCCCAATAATCAAAA | 30 |
| 126 | TGAAGCCTTTACAAAAACGTCAAAGTAATTGA | 32 |
| 127 | CAGTACAACGCCCACGCGTTGAAAGGCACCAA | 32 |
| 128 | GGTTTTTCGCCCTTCAAGCCCGAGATAGGGTT | 32 |
| 129 | TAATCTTGTCATCAGTTGGGAACAGAAAAC | 30 |
| 130 | CCCCACCAGAGCCACCCGTCACCACCCC | 28 |
| 131 | GCTTATCTTGTTTATAAACAGCAAGAAA | 28 |
| 132 | GTGCCGTACCGATTTACTGCGCGTCTACAGGG | 32 |
| 133 | CCTCATTTGTTTTAACGCTGAGACGCCAGCAT | 32 |
| 134 | AACGTGCTGAGTAGACCGTTGTACATTCT | 29 |
| 135 | TCCAATAAATCAATAAACAATAACGGATTC | 30 |
| 136 | AGGCACCGACAATCATATGCGTTATACAAATCCCC | 35 |
| 137 | GGCCAACATAAAACATCGGTCAGTTCAATATC | 32 |
| 138 | CAATGAAAGGAATACCAGAAAATAGCGCCAAA | 32 |
| 139 | GGGAGCCCAAGCACTATTGGAACAAGAGTCCA | 32 |
| 140 | CAACTGTTTCACAATTGTAATCATACGCGCGG | 32 |
| 141 | AGCGTAAGCTATTAGTACGCTGAGACCTTGCT | 32 |
| 142 | GGTGAGAAATGTGTAGAGGCAAGGCATTAACA | 32 |
| 143 | GGTAATAATCAGGGATTGCCGTCGGCGGAGTG | 32 |
| 144 | AATGGGATTTGTTAAATTTAAATTATCTACAA | 32 |
| 145 | TGATAATCCTCAGGAATAACAACCTCACGACG | 32 |
| 146 | CACCCGCCGAGCTAAAGTGAGACCCCTAAA | 30 |
| 147 | ACAACCATACTACAACCAGTTTCAAGAGGGTT | 32 |
| 148 | TTGAATGGAATACGTGAGGAAAAAATATTACC | 32 |
| 149 | GAGTTGCAGGTTTGCGTTTCCAGTCATACGAG | 32 |
| 150 | TGAAAGAGTTTTCATTTATCCTGTACACTA | 30 |
| 151 | CCCCGTAGATGGGCGCATGCGGGCCTCTTCGCTATTCCCC | 40 |
| 152 | TACTAGAACCTCCGGCAACATAGCTAATTTTC | 32 |
| 153 | AGAAAGATACAAGAAGCTTGCCGAGATTTG | 30 |
| 154 | CCAAGTACCCATATTTCGACGACAATCATAAT | 32 |
| 155 | CCCCCCGATTGAGGGACTGGCATGATTAAGACTCCCCCC | 39 |
| 156 | GCAAATGAATGTCAACCAGCTTTAATATTT | 30 |
| 157 | TGGTCAGAAAGAAAGTTATTATTTCAGG | 28 |
| 158 | CCTTTTTTCAGATGAAAATGGAAGAGGAGCGG | 32 |
| 159 | CAAGCGCGGTAATGCCCACCCTCACAATGACA | 32 |
| 160 | TTTGCCAGTAAATCAAAAATCAGAGTATTAAA | 32 |
| 161 | AGAGAATAAGAGCCATATTATTTATCCCAATCCCC | 35 |
| 162 | CCCCGCGATGGCCCACTACGTGAA | 24 |
| 163 | TAAAATACAAACAAAGGAACGAGGATCAAGAG | 32 |
| 164 | CCCCCATTATCATTTTGCGGAACTTGG | 27 |
| 165 | GCTTTCCGTTGCAACGCACAGACCATCAAC | 30 |
| 166 | AAATATTGTAGCAAGGGCAGCACCCCATCTTT | 32 |
| 167 | CCCCCTATTTTTGAGAGGTAAACGTTAATATTTTGTCCCC | 40 |
| 168 | CTTTTACATCGGGAGATAATCCTGCAGATGAT | 32 |
| 169 | CCCCCCGCCACCCTCAGACGATCTAAAGTTTTGTCGCCCC | 40 |
| 170 | GATGGTTTATGCGATTCAGATACATTGCAAAA | 32 |
| 171 | TAGCAGCCTAGCAAGCGATTAGTTAAGAAAAA | 32 |

| | | |
|---|---|---|
| 172 | CCCCATGAATCGGCCAGGTCATAGCTGTTTCCTGCCCC | 38 |
| 173 | CCCCATGAAACCGGCGACATTCAACCCC | 28 |
| 174 | CCTTTTTACCAGAAGGAAAGAAACCACAATCA | 32 |
| 175 | CCCCTTATAGTCAGAAGAACTAAAGTACGGTGTCTGCCCC | 40 |
| 176 | CAGGGAAGAGGCTTGCAAAAGAAAATCTTA | 30 |
| 177 | GCGCTAACAAACGTCAAAAGAAGGGAAGGT | 30 |
| 178 | CCGTACTCTTTCGGAATAAACAGTTAATGCCCCCCC | 36 |
| 179 | TACCGAGCGCCAGGGTCGCCATTCCGACGACA | 32 |
| 180 | CCATCACGATTAAAGGTTTGACGAGCGCTGGC | 32 |
| 181 | CCCCAATAACCTTGCTTCTGTAATATC | 27 |
| 182 | ATTGCATCGGATAGCAAACAGTTGAAAAATC | 31 |
| 183 | GAATTAGACGTCACCGTATTTTGTGCAAAGAC | 32 |
| 184 | CCCCTTGAGTTAAGCCCAATAGATAACCCACAAGAACCCC | 40 |
| 185 | CCCCTAAAATTCGCATTAAATTTAGGTCACGTTGGTCCCC | 40 |
| 186 | CCCCCGAGGTGAATTTCCAACGGCTACAGAGGCTTTCCCC | 40 |
| 187 | TAGATAATCAAACAATCTGATTATATTGTTTG | 32 |
| 188 | GACTGTAGTTTTGATGCGGGGTTCATTTGG | 30 |
| 189 | TGACAGGAGGTTGAGGCCAGAATGTGCCTTGA | 32 |
| 190 | GCAAACAATTAAGCAATATTTTAATACAAAAT | 32 |
| 191 | TTTAGACTAAAAAGATTAAGAGGAATGCTGT | 31 |
| 192 | CAGGCAAAATAAAGTGCTAGAGGAGCCAGGGT | 32 |
| 193 | AATATATTTCATCTTCGGGTAATATAGGAATA | 32 |
| 194 | GGTCGCTGCGCATTAAAGCAGATAATTGCG | 30 |
| 195 | CATGTTACTCGGAACGTTTCCATTTAATTGT | 31 |
| 196 | GCGAAAAGGAGCGGAGAAAGGAGCTAAACA | 30 |
| 197 | TTGTAAAAGGTCGACTTAAAGCCTAATTGCGT | 32 |
| 198 | GATATAAGATTAAGAGGGGGTCAGGAAAGCGC | 32 |
| 199 | CCGGAAGCGCGCCATTTTTCCCAGCGTCGGAT | 32 |
| 200 | TCTCCGTGGAACGCCACCCAAAAAAGTCTGGA | 32 |
| 201 | CCCCCAGAAATAAAGAAATTTTATTTGCACGTAAAACCCC | 40 |
| 202 | AGACAGCATTAGCCGTACAACGCTGACGAG | 30 |
| 203 | GATAGCCCGAGATAGAGACGCTCAAACTATCG | 32 |
| 204 | TATAAATCCTGCCCGCTATTGGGCTCCCCGGG | 32 |
| 205 | CCCCTCTTACCAGTATAAAGCCAGACG | 27 |
| 206 | ATCGGTTTATGAATTTTTAGCGTAAACCGCCA | 32 |
| 207 | ACCAATAGGGAACAAATGAGGGGAAGGCTGCG | 32 |
| 208 | AGCTGATTTTTTCACCCTCACATTGGGGTGCC | 32 |
| 209 | AGCACTAAGCCCGAACCCACCAGAGGTTAGAA | 32 |
| 210 | CCCCCCTGCCTAAGGAGGTTTAGTACCCC | 29 |
| 211 | TCAATATGCCCTCATATAAAGCCTAAAGGTGG | 32 |
| 212 | CCCCGTAATAACATCACTTGCCTTTCCTCGTTAGACCCC | 39 |
| 213 | TTGTATAAGCCAGTTCGGCGGAGATTAAGT | 30 |
| 214 | ACAGGTCAACCCTCGTACACCGGAATAAACAA | 32 |
| 215 | CCCCGAAGTTTCATTCCATATAAATTTCGCAAATGGCCCC | 40 |
| 216 | CTTTTTAAAAAGCCTGACAGTAGGCAACATGT | 32 |
| 217 | AGACTTTAACATTTGAAAAGCATCAGCCAGCA | 32 |
| 218 | TTTAACGTAATGGAAACTATTAATGATAGCTT | 32 |
| 219 | GCGCCCAATTTACAGATCTTTCCAACCGAAGC | 32 |
| 220 | AACTATATGTAAATGCCCGGAAGCTTAATTGC | 32 |
| 221 | CCCCACGCCAGCTGGCGAAAGGGGGATGTGCGCGATCGGTCGTAACC | 47 |
| 222 | ACCAGAACCACCACCAGTTCCAGTAGTGTACT | 32 |
| 223 | CGTTAAATAAGAATAATTACCAGAGAGCAACA | 32 |
| 224 | AAGATGATGAAACAAAGTCATTTTTTCGAGCT | 32 |
| 225 | ATTAATTACATTTAACGAGTACCTAAACTCCA | 32 |
| 226 | GTTTGAAATACCGACCAGAGGCTTTAACGCCA | 32 |

*Table S3: List of staples with surface binding function.*

| Staple number | Sequence (5' to 3') | Nucleotide length |
|---|---|---|
| 1 | Biotin-TTCCATCACCACGTGGCGGCGCTAGGGCACGTAT | 34 |
| 2 | Biotin-TTATGAAAGTTATAGCCCCCCTCAGACATTCCAC | 34 |
| 3 | Biotin-TTGCCTGATTGCTTTGAAAGTAGTAGCAAAGAAT | 34 |
| 4 | Biotin-TTCGCCTCCCTCAGAGCCAGCGTTTGGTAATCAG | 34 |
| 5 | Biotin-TTATTACCTTAATTTCAAGAACGGTGACCAACTT | 34 |
| 6 | Biotin-TTGGAGAGGCGCAAGCGGAATCGGCAAAATCCCT | 34 |

*Table S4: List of staples with fixed fluorophores.*

| Staple number | Sequence (5' to 3') | Nucleotide length |
|---|---|---|
| 1 | TCAAAGCGAACCAGATGATGCAAATCCAATTT-ATTO 542 | 32 |
| 2 | TTAGATACCAGTTGATAAATATGCCAAAGCGGTTT-ATTO 647N | 35 |

#### S12. 8HB DNA origami staples list

*Table S5: List of staples for the origami core.*

| Staple number | Sequence (5' to 3') | Nucleotide length |
|---|---|---|
| 1 | GGGCGAAACAATCAATTGAGGATTGAACGTTA | 32 |
| 2 | ACTATTAGCCAAAGATAGATTTTATCAT | 28 |
| 3 | TTGAGTGTTGAGGGAGATCTAAAAAAGGAGCG | 32 |
| 4 | TTATAAAATTCATTAAATCAACTCAGATG | 29 |
| 5 | TTTGATGGACTTGAGCTCAAACCCGATTGTTT | 32 |
| 6 | CGCTGGTATCACCATCTAAAGGGGTTAG | 28 |
| 7 | CTGGCCCTCCGGAAACTGCCACGCCACGTAAA | 32 |
| 8 | AGACGGGAGCACCGAGGCGGTTTTTCAG | 28 |
| 9 | GGGCGCCATTGCCTTTACGAACCATAACAGTA | 32 |
| 10 | CCAACGCATCGGCAAGCCCTAACGGATT | 28 |
| 11 | AACCTGTCGCGTTTGCAATGGCTATTACAAAA | 32 |
| 12 | GTTGCGCGGAACCAGTAAGAATCAATTA | 28 |
| 13 | GCCTAATGTCAGAGCCCAACAGAGCAAACATC | 32 |
| 14 | AGCCGGAAGAGCCAACACGACAACAATT | 28 |
| 15 | TGTTATCCCCACCACCATGGATTAGAAACAGT | 32 |
| 16 | ATCATGGAGGTTGAGAGTAATGCGAGAG | 28 |
| 17 | GAGGATCCTATTCACATAGAACCCGAGGGGGT | 32 |
| 18 | GCCAAGCAGAATGGGCCTTTAAGCGTCC | 28 |
| 19 | CCCAGTCATTCCAGTACATTATGATTCATTGA | 32 |
| 20 | AGGCGATTACAGGAGAGCATATTCAGAA | 28 |
| 21 | TACGCCAGGGGTCAGTAAGAATTAATCAGGTC | 32 |
| 22 | GAAGGGCAACAGTTTTAACATAGCAAAG | 28 |
| 23 | GCCATTCGCTATTATTAGGTGGCAGGAAGCCC | 32 |
| 24 | ACCGCTTCTGAGACGCTATATAATTCGA | 28 |
| 25 | CAGGAAGAGCGGGGTTATACATTTGCAAACTC | 32 |
| 26 | CAGTTTGGCCGTCGTGCGAACCTTTAAT | 28 |

| | | |
|---|---|---|
| 27 | TGTAGATGGGAATAGGTTTCATTCTTTGCGGA | 32 |
| 28 | CGCGTAACCACCACACGGAAGGTTGGAAGGTAGAACTGGC | 40 |
| 29 | GCCGCTACAGGGCGCCAGTTGGCAAAGGTGAAGGAAACG | 39 |
| 30 | AATCAGAGCGGGAGCATGAAAAAGTAGCACCTTACCGAA | 39 |
| 31 | TAAAGGGATTTTAGACTGCAACAGGTCACCAAGAGCAAGA | 40 |
| 32 | AATCCTGAGAAGTGTCAGAGGTGTAATCAGTCCCACAAG | 39 |
| 33 | GCCACCGAGTAAAAGAAATACCGAAGCGTCAGAGGGTAAT | 40 |
| 34 | AGTAATAACATCACTTTATTTTTGCATCTTTTTAACATAA | 40 |
| 35 | TCAAACTATCGGCCTCTGAAAGCGAGCCACCAAAATGAA | 39 |
| 36 | AACAATATTACCGCCAATTCTGGCGCCACCCTCCAAATAA | 40 |
| 37 | AAACGCTCATGGAAACACCAGTCCCACCCTCCAAAATAA | 39 |
| 38 | TCAAATCACCATCAACAATGCCTGGCAGGTCTGATACCG | 39 |
| 39 | TCTAGCTGATAAATTAAAAATTTTAACAAATAGCTTGCTT | 40 |
| 40 | CTATTTTTGAGAGATCGGGAGAAAAAGCGCAAAAAGGAG | 39 |
| 41 | GTCATTGCCTGAGAGTACCAAAAAAAGCGTCATTTCACGTT | 40 |
| 42 | GTCAATCATATGTACCGCAAGGCAGCCTTGAGTCAACAGT | 40 |
| 43 | AAAGCCCCAAAAACATAGTAGCAAATGCCCCATTTTCTG | 39 |
| 44 | AAATATTTAAATTGTAAGCTGAAACTGAAACAGTTTTGTC | 40 |
| 45 | TAAAATTCGCATTAACCTGTTTATCCTCAAGACAGCCCT | 39 |
| 46 | GGAATTGACCGCCGCGATAGCCCGAGATAGGG | 32 |
| 47 | TATCTGGTGTACTATGAAATCGGCAAAATCCC | 32 |
| 48 | AGCAGCAATAAACAGGTGCAGCAAGCGGTCCA | 32 |
| 49 | ACACCGCCAGGAACGGGATTGCCCTTCACCGC | 32 |
| 50 | AGATAAAATTTTATAATTTCTTTTCACCAGTG | 32 |
| 51 | CCATTAAAGTCTGTCCAGGCGGTTTGCGTATT | 32 |
| 52 | ACAGACAAGCCTGAGTCGCTTTCCAGTCGGGA | 32 |
| 53 | CTTCTGACTGCTGGTAAACTCACATTAATTGC | 32 |
| 54 | AAAGGGACGCCATTGCGTGTAAAGCCTGGGGT | 32 |
| 55 | GGCAGATTTACCTACATTCCACACAACATACG | 32 |
| 56 | TTTAAATGTATGATATCGAGCTCGAATTCGTA | 32 |
| 57 | CAAGGATAATGCCGGACTGCAGGTCGACTCTA | 32 |
| 58 | TACTTTTGCTACAAAGTAAAACGACGGCCAGT | 32 |
| 59 | TCGGTTGTCTGGAGCAGTAACGCCAGGGTTTT | 32 |
| 60 | TCATACAGCCGGTTGACGGGCCTCTTCGCTAT | 32 |
| 61 | ACTAATAGGGAAGATTGCTGCGCAACTGTTGG | 32 |
| 62 | GGGGCGCGAACGTTAAGGAAACCAGGCAAAGC | 32 |
| 63 | GTCAATAAATTTTTGTCAGCCAGCTTTCCGGC | 32 |
| 64 | ATGGCAATACCGGAATGTCCTGAACAAGAAAA | 32 |
| 65 | AACCTACCACCTAAATCGGCTGTCTTTCCTTA | 32 |
| 66 | GTTTAACGAATCGCAAAAGCAAGCCGTTTTTA | 32 |
| 67 | CCTTTTACATATAACTCATTACCGCGCCCAAT | 32 |
| 68 | TCGCGCAGGTGAATTTGAGGCGTTTTAGCGAA | 32 |
| 69 | AAGAAAACTTTCCCTTCTATTTTGCACCCAGC | 32 |
| 70 | TCATTTGATTGCTTCTAATCTTACCAACGCTA | 32 |
| 71 | GCTTTTGCAGTAAGAGCAGGGAGTTAAAGGCC | 32 |
| 72 | AATACTGCAGATTTAGGAGGGTAGCAACGGCT | 32 |
| 73 | ATCCCCCTACATTATTCTAAAGACTTTTTCAT | 32 |
| 74 | TTTACCCTTCATTATAGCACCAACCTAAAACG | 32 |
| 75 | GAAAGACTTTGAGATGATTATACCAAGCGCGA | 32 |
| 76 | GCTTCAAAGAGAAACAAGATTTGTATCATCGC | 32 |
| 77 | ACTAGAAAAAGCCTGTATCATATTACCCAAAAAATATTGA | 40 |
| 78 | TTAAATAAGAATAAACTCATCAATGGAAACCGATTATCAC | 40 |
| 79 | TTAATTTCATCTTCTGATATCAAATAGCTATCATTACCAT | 40 |
| 80 | AACGCGAGAAAACTTTAAAGAAATAATAATAATGAAACCA | 40 |
| 81 | ATGCTGATGCAAATCCTCAGATGAAGAGATAAAGCGACAG | 40 |

| | | |
|---|---|---|
| 82 | CGGCTTAGGTTGGGTTATCGGGAGAAAGTCAGACTGTAGC | 40 |
| 83 | CTGAGAAGAGTCAATAAGGCGAATAGAGAGAACATAATCA | 40 |
| 84 | TGAAAACATAGCGATAAAAGAAGATAACGTCAACCGGAAC | 40 |
| 85 | GTCGCTATTAATTAATAAAATTAAATCCCAATCAGAACCG | 40 |
| 86 | ATGTGAGTGAATAACCATTACCTTGCCAGTTAAGAGCCGC | 40 |
| 87 | AGGAATTACGAGGCATAAAAGAAGAAACAGCTAGACGATT | 40 |
| 88 | ACATTCAACTAATGCAAATGTTTAGTTTATCAAATCCTCA | 40 |
| 89 | AAAGATTCATCAGTTGGGAATCGTAAGGCTCCGTCTCTGA | 40 |
| 90 | ACGAACTAACGGAACACAAATGCTATAATTTTACATGGCT | 40 |
| 91 | GATTTTAAGAACTGGCGACTATTAAACAACTTTAACAGTG | 40 |
| 92 | TCAACTTTAATCATTGATCAAAAAGTAAATGACTGCCTAT | 40 |
| 93 | GAGTAGTAAATTGGGCTCAAATATGATCTAAATGAAAGTA | 40 |
| 94 | TAAGGCTTGCCCTGACGCGAACCATTCCACAGAGAAGGAT | 40 |
| 95 | AGTAATTCTAGAAAATGAGTAACAAGAGCCGT | 32 |
| 96 | ACATGTTCGACTCCTTACCACCAGTATCTTTA | 32 |
| 97 | ATCAACAAAACGGAATCCTGATTAAGTTGAAA | 32 |
| 98 | ATAATATCTACCAGAAATAATCCTTCAATCAA | 32 |
| 99 | AAACCAATTTAAGAAATAATGGAACATCACCT | 32 |
| 100 | TCATTCCAAATAGCAAATTATTTGTGAGAGCC | 32 |
| 101 | GCACTCATTTAAGCCCTGCGTAGACAGTATTA | 32 |
| 102 | TTTTCATCAATATCAGATATACAGCCAGCAGA | 32 |
| 103 | AGCAAGCACCCTGAACAAACAATAAAACATCG | 32 |
| 104 | GGTATTCTAAGCGCATATACCAAGTTAGTCTT | 32 |
| 105 | CCTCCCGAGCCTTTACTATTCATTTACGTGGC | 32 |
| 106 | AAATCAAGTTTTTGTTTGATGAAAATAGAACC | 32 |
| 107 | TACAATTTTATTATTTTTACATTTCAGTAATA | 32 |
| 108 | ACGAGCGTCTTTCACGATCAATAT | 24 |
| 109 | GGTCGCTGCGCCGACACCAAAATAGTGTAGGT | 32 |
| 110 | GCTTTTGCAATTTCTTTTTTGCCATCATATAT | 32 |
| 111 | AGACAGCATTGTATCGGACTGGATTTTCAACG | 32 |
| 112 | ACAGAGGCCCAAAAAACATAAATACCCTGTAA | 32 |
| 113 | GAGGAAGTTGCGAATATTAAACAGAAGCTAAA | 32 |
| 114 | TAATGCCAAGTGAGAAAATCAAAAGCAAAATT | 32 |
| 115 | AAAGAGGCTTTTGCTATAGTCAGACCAATAAA | 32 |
| 116 | TCTTTGACAGACGTTAGATTAAGATCAATTCT | 32 |
| 117 | AACAAAGTAGCGTAACCGCGTTTTTTTCATTT | 32 |
| 118 | CTGATAAACTGTAGCAGACCGGAACGCAAATG | 32 |
| 119 | CCATGTTACTGAGTTTGAGAGTACGAGTAGAT | 32 |
| 120 | GGTCAATCCAAGCCCAAGGTCATTCATATAAC | 32 |
| 121 | TTGAAAGACTCAGAGCTGCTGAATCAACTAAA | 32 |
| 122 | CAACATATATATAAAGATCGCCATATTTAACA | 32 |
| 123 | GCAAACGTGTCCAGGCCAACGCTCAACAG | 29 |
| 124 | ATGATTAAAGCTAATGATATGCGT | 24 |
| 125 | CAATAATTAGATAACATAATT | 21 |
| 126 | AACAAAGTCCATCCTAATAAGGCG | 24 |
| 127 | GCCCTTTCAATAATTTAATGGTTTGAAAT | 29 |
| 128 | AACAATGAAGAACGGGTATTTTAG | 24 |
| 129 | AATTGAGCGAGAACGACAAAG | 21 |
| 130 | TGAGCGCTGTAGGAATATATGTAA | 24 |
| 131 | CTGAACAAATCAGATAACCTC | 21 |
| 132 | AAACAGGGAAGAACGCATCAAAATCATAGGTC | 32 |
| 133 | AATAGCACTTGCGGTAAGACG | 21 |
| 134 | GAAACGATATTAGTTGAGAATCCT | 24 |
| 135 | ACAGCCATATCCTGGTAAATC | 21 |
| 136 | CCCCAGAGCCTAATTTTTTTAATGTTTACATT | 32 |

| | | |
|---|---|---|
| 137 | ATAGTTGAGGCTTGCAACACTATCATAAC | 29 |
| 138 | TCGAGGTGGGGATCGTACGCCAAA | 24 |
| 139 | CCTTTAATCGGAACGAATACC | 21 |
| 140 | GAAAATCTTTTGAGGAACAGGTAG | 24 |
| 141 | AAGGAATTTCCATTTAATAAA | 21 |
| 142 | TTCAGCGGCTACGAAGCCAGTCAGGACGTTGG | 32 |
| 143 | TATGGGAAAAAGAACTTATGC | 21 |
| 144 | GTCTTTCCCCCCAGCGGTTTAATT | 24 |
| 145 | CATAGTTACAACGGCCAGAAC | 21 |
| 146 | TACAACGCTTGTGTCGTCAGTGAA | 24 |
| 147 | CGTAACACTTAGCCTTACCCAAATCAACG | 29 |
| 148 | AGGGATAGATAAGGGACAAGAGTA | 24 |
| 149 | TTTTGTCAAACCGTCTTGGCGAGAAAGGAAGG | 32 |
| 150 | TTACCAGCAAGAACGTGCGGGCGC | 24 |
| 151 | CAACCGATTGTTCCAGTCACGCTG | 24 |
| 152 | CGGAAATTTCAAAAGACTTAATGC | 24 |
| 153 | CGTCACCGTGGTTCCGGTTGCTTTGACGAGCA | 32 |
| 154 | CCAGCAAATTGCCCCACTCGTTAG | 24 |
| 155 | TAGCAAGGGAGAGAGTAGGCCGAT | 24 |
| 156 | TCGATAGCCAACAGCTTACGCCAG | 24 |
| 157 | AATCAAGTGGGTGGTTTCAGTGAG | 24 |
| 158 | GCGTTTTCGCGGGGAGATCACGCAAATTAAC | 31 |
| 159 | CCTTATTAGTGCCAGCCTTTGATT | 24 |
| 160 | AAATCACCTCACTGCCAGAAGAAC | 24 |
| 161 | CGCCTCCCAGTGAGCTATATCCAG | 24 |
| 162 | CCACCCTCAGCATAAAAACAGGAA | 24 |
| 163 | CACCAGAAGCTCACAATTTTGACGCTCAATCG | 32 |
| 164 | TTGACAGGTCATAGCTGGAGACAG | 24 |
| 165 | GGCCTTGACCGGGTACTCAACCGT | 24 |
| 166 | TTAAAGCCTTGCATGCGAGGGTAG | 24 |
| 167 | ATTTACCGCGACGTTGGCTATCAG | 24 |
| 168 | TTTGATGATAAGTTGGAACAAGAGAATCGAT | 31 |
| 169 | TTTTAACGCTGGCGAAACTAGCAT | 24 |
| 170 | CCCGTATAGATCGGTGTAATCAGA | 24 |
| 171 | TTCGGAACCCATTCAGGTATAAGC | 24 |
| 172 | TTAAGAGGCTGGTGCCTATTTTGT | 24 |
| 173 | TAGGATTATCGCACTCTAAATCAGCTCATTTT | 32 |
| 174 | GGATAAGTAGGGGACGCCATCAAA | 24 |
| 175 | TATAGCCCGGCGCATCGTAGCCAG | 24 |
| 176 | TTGAAGTACCGCTAAATATG | 20 |
| 177 | CCCGGAACAAACGGCGGA | 18 |
| 178 | CCCACATGTTTCACCCTCAGAACCCCC | 27 |
| 179 | ATAATGCTGTAGCTCACCC | 19 |
| 180 | ACGCCAACATTAAATCCGCAAAGACACCACCC | 32 |
| 181 | CCCGTGTACAGACCAG | 16 |
| 182 | CCGCCACCGGACAGATGAACGCCC | 24 |
| 183 | CCCGCCACCCTCAGAA | 16 |
| 184 | CGTATAACCTCAAATACATTTGGGGATAGCCG | 32 |
| 185 | TGCTGAACGTGCTTTCGCAGGCGAAAATCCTG | 32 |
| 186 | GGATTATATGTGATAAATTTACGAGCATGTAG | 32 |
| 187 | ACCGACCGCTTCTGAAAGTAAGCAAATTAGAG | 32 |
| 188 | CGTTGTAGGAACTGATTTTTCGGTAGAATTAA | 32 |
| 189 | TAATGCGCCAATACTTTGCATTAATGAATCGG | 32 |
| 190 | CGCCTGATTACCTTTTTATAGAAGGCTTATCC | 32 |
| 191 | TGAGAGACTGCTTTGATAGACGGGCATAGCCC | 32 |

| | | |
|---|---|---|
| 192 | TCTGGTGAGAAAGGCCGTTTCCTGTGTGAAAT | 32 |
| 193 | AAAGATTCAAAAGGAAAGAGCCGCAACCATCG | 32 |
| 194 | ACAACGACGATAAAAAATGACAACCGCCAGCA | 32 |
| 195 | CCTCGTTTACCAGTAACATAACCGATATATTC | 32 |
| 196 | GAACGGTAAAGCCTCAGTGTACTGAACAACTA | 32 |
| 197 | AAGCAATAATCGTAAAAGGGGGATGTGCTGCA | 32 |
| 198 | AACGAGAAATCTACGTAAACGGGTAAAATACG | 32 |
| 199 | GAAGAAAATGACCATATAGAAAGGGTAATAAG | 32 |

*Table S6: List of staples with DNA-PAINT binding strands.*

| Staple number | Sequence (5' to 3') | Nucleotide length |
|---|---|---|
| 1 | CCCGGAAAGCCTACAAACAATT TTATCTCCTATACAACTTCC | 22+20 |
| 2 | TTAGACTTGGCGAACGATCAAGAGCTTGACGG TTATCTCCTATACAACTTCC | 32+20 |
| 3 | CCCCGACAACTCGTATGTAATTTAGGC TTATCTCCTATACAACTTCC | 28+20 |
| 4 | CCCCGGAATAAGTTTA TTATCTCCTATACAACTTCC | 16+20 |
| 5 | TATACAAATTCTTACCACAAAGAAATTACGCAGCGACATT TTATCTCCTATACAACTTCC | 40+20 |
| 6 | TTTGCGGAAGTATAAAACGACGACAATAAACA TTATCTCCTATACAACTTCC | 32+20 |
| 7 | GGAGCACTTGTAGCGGTTTGGAACAAGAGTCC TTATCTCCTATACAACTTCC | 32+20 |
| 8 | TAGGGCGCTGGCAAGAACAACTAACAAAAGGGTATGTTA TTATCTCCTATACAACTTCC | 39+20 |
| 9 | GAAGAAAGAATACATTAGAAAATTAAAGGTGG TTATCTCCTATACAACTTCC | 32+20 |
| 10 | CAATAGATCGAAAGGAGGACTCCAACGTCAAA TTATCTCCTATACAACTTCC | 32+20 |
| 11 | TTAATTTTAATTGAGATACCGACAAAAGGTAA TTATCTCCTATACAACTTCC | 32+20 |
| 12 | TAGGGCTTAAAAGTTTACATACATCATATGGT TTATCTCCTATACAACTTCC | 32+20 |
| 13 | ATCTTGACAAGAACCGTTGATAAGATAGGAACGATATAAG TTATCTCCTATACAACTTCC | 40+20 |
| 14 | GCGCATAGGCTGGCTGAGCTTAATCACCACCCCGTACTCA TTATCTCCTATACAACTTCC | 40+20 |
| 15 | TGCTCCTTGATATTCAGGAACGAGGCGCAGAC TTATCTCCTATACAACTTCC | 32+20 |
| 16 | AGTTGATTGCCTTCCTGTAACCGTGCATCTGC TTATCTCCTATACAACTTCC | 32+20 |
| 17 | GTACGGTGATGTGAGCGGATAGGTCACGTTGG TTATCTCCTATACAACTTCC | 32+20 |
| 18 | AATAATTCGCGTCTGCCCAATTCAGAGGGTTCCATGTAC TTATCTCCTATACAACTTCC | 39+20 |
| 19 | CTTTCATCAACATTAATCTGGAAGTGTATCACTCATTTTC TTATCTCCTATACAACTTCC | 39+20 |
| 20 | TTAACCAAACCATTAGTTGCTCAGGTACAAAC TTATCTCCTATACAACTTCC | 32+20 |
| 21 | TTAGTTTGTAGGAACGACGACAGTATCGGCCT TTATCTCCTATACAACTTCC | 32+20 |
| 22 | CAACAGGTCTGCTCATAAATCCGCGACCTGCT TTATCTCCTATACAACTTCC | 32+20 |
| 23 | TAACAAAGCAGGATTACGTCACCATACCAGGC TTATCTCCTATACAACTTCC | 32+20 |

*Table S7: Imager strand sequence.*

| Imager number | Sequence (5' to 3') | Nucleotide length |
|---|---|---|
| 1 | BHQ2-AAGTTGTAATGAAGA-Cy3b | 15 |

*Table S8: List of staples with surface binding function.*

| Staple number | Sequence (5' to 3') | Nucleotide length |
|---|---|---|
| 1 | Biotin-GAATTATCTTAGTATCCAGAACGCGCCTGTTT | 32 |
| 2 | Biotin-ACAGAAATTTCAAATATATTAAACCAAGTACC | 32 |
| 3 | Biotin-CCTGAGCAGCTTAGATGAGGTTTTGAAGCCTT | 32 |
| 4 | Biotin-AATAGTAAGATACATACACCCTCAGCAGCGAA | 32 |
| 5 | Biotin-CGGATTGCTGAATTACTACACTAAAACACTCA | 32 |
| 6 | Biotin-TGGCTTAGACCTTCATACCGAACTGACCAACT | 32 |

*Table S9: List of staples with fixed fluorophores.*

| Staple number | Sequence (5' to 3') | Nucleotide length |
|---|---|---|
| 1 | GGAGGTTTCCGTAATGGAGTAACAACCCGTCGGATTCTCCGTGCCC-ATTO 542 | 46 |
| 2 | ATTO 542-CCCAGAGGCATTTTCGAGCCAGTAATAAGAGAAAAAGAAACTTTGCCCTAGA AGTA | 56 |

#### S13. Experimental parameters of recorded videos

*Table S10: ISO parameter is reported for smartphone cameras. Irradiance is reported as the equivalent top-hat excitation using the reported waist (see Methods) and the measured power at the sample plane.*

| | $\lambda_{exc}$ (nm) | Irradiance ($kW/cm^2$) | Exposure time (ms) | Number of frames | Sample, DNA origami, and fluorophore | Imaging buffer | ISO | Setup | Comments |
|---|---|---|---|---|---|---|---|---|---|
| Figure 2b | 640 | 0.68 | 100 | - | 2LS with ATTO 647N | PPC | - | High-end microscope | Single-molecule intensity traces. 10 frames average |
| Figure 2c | 532 | 0.76 | 250 | - | 2LS with ATTO 542 | PPC | 1000 | Smartphone-based with Samsung Galaxy S22 Ultra | Single-molecule intensity traces. 16 frames average |
| Figure 3b | 532 | 0.76 | 250 | 40785 | 8HR with Cy3b | PPC | 1000 | Smartphone-based with Samsung Galaxy S22 Ultra | DNA-PAINT video. 3 frames average for analysis. 750 ms effective exposure time |
| Figure 3c | 532 | 3.2 | 100 | 18000 | 8HR with Cy3b | PPC | - | High-end microscope | DNA-PAINT video |
| Figure 4a | 532 | 0.76 | 200 | 28245 | Microtubules (U2OS cell) targeted with Cy3b | PPC | 640 | Smartphone-based with Samsung Galaxy S22 Ultra | DNA-PAINT video. 3 frames average for analysis. 600 ms effective exposure time |
| Figure 4d | 532 | 1.9 | 50 | 36000 | Microtubules (U2OS cell) targeted with Cy3b | 1xPBS | - | High-end microscope | DNA-PAINT video |
| Figure S5a | 532 | 0.76 | 250 | - | 2LS with ATTO 542 | PPC | 600 | Smartphone-based with iPhone 14 Pro | Single-molecule intensity traces. 16 frames average |
| Figure S5b | 640 | 1.8 | 100 | - | 2LS with ATTO 647N | PPC | - | High-end microscope | Single-molecule intensity traces. 10 frames average |
| Figure S6a | 532 | 0.76 | 80 | - | 2LS with ATTO 542 | PPC | 2850 | Smartphone-based with Huawei P20 Pro | Single-molecule intensity traces. 48 frames average |
| Figure S6b | 640 | 0.68 | 100 | - | 2LS with ATTO 647N | PPC | - | High-end microscope | Single-molecule intensity traces. 10 frames average. |

| | | | | | | | | | |
|---|---|---|---|---|---|---|---|---|---|
| Figure S8 | 532 | 0.76 | 250 | 40785 | 8HR with Cy3b | PPC | 1000 | Smartphone-based with Samsung Galaxy S22 Ultra | DNA-PAINT video. 3 frames average for analysis. 750 ms effective exposure time |
| Figure S9 | 532 | 3.2 | 100 | 18000 | 8HR with Cy3b | PPT and PPC | - | High-end microscope | Buffer performance. DNA-PAINT video. |
| Figure S12 | 532 | 0.76 | 250 | 51099 | 8HR with Cy3b | PPC | 1000 | Smartphone-based with Samsung Galaxy S22 Ultra | DNA-PAINT video. 3 frames average for analysis. 750 ms effective exposure time |
